# The human papillomavirus L2 protein in the capsid is an ensemble of related structures poised to exit the virus particle

**DOI:** 10.64898/2026.08.18.745536

**Authors:** Changin Oh, Huaxin Yu, Patrick M. Buckley, Frank L. Horrigan, Kashif Mehmood, Jeongjoon Choi, Jun Liu, Daniel DiMaio

**Affiliations:** Department of Genetics; Department of Microbial Pathogenesis; Department of Molecular Biophysics and Biochemistry; Department of Therapeutic Radiology; Yale Cancer Center

## Abstract

The human papillomavirus (HPV) capsid contains 360 molecules of the L1 protein, which assembles into pentamers that form the capsid shell, and up to 72 molecules of the L2 protein. The C-terminus of the L2 protein must emerge from the capsid, insert into the endosome membrane, and protrude into the cytoplasm to bind to cellular proteins that mediate trafficking of the virus to the nucleus during virus entry. Computational analysis predicts that much of L2 is disordered, but the stoichiometry, structure and location of L2 in the capsid remain elusive. Here, we use cryo-EM single particle analysis of HPV16 pseudovirus capsids with and without the L2 protein combined with AlphaFold3 predictions and molecular dynamics simulations to develop a structural model of L2 within the capsid. Our results revealed that a single molecule of L2 associates with the inner surface of most L1 pentamers. The relatively rigid structure of the pentamer forces the long intrinsically disordered segments of L2 to adopt numerous related, compact, stable conformations in the capsid, whereas the structured elements of L2 remain folded but adopt varied positions in space largely beneath the pentamer. A finger-like projection of L2 extends into the central pore of the pentamer, but many different segments of the disordered domains of L2 can form fingers. In some models the C-terminus of L2 extends through the pore to the exterior surface of the capsid positioned to insert into the endosomal membrane. Consistent with these models, analysis of purified pseudoviruses showed that the C-terminus of multiple L2 molecules is exposed on the surface of each capsid. These findings provide a structural basis for understanding how L2 is organized in the capsid poised to initiate its action during virus entry.

## INTRODUCTION

Human papillomaviruses (HPVs) trigger approximately 5% of cancer. Despite their medical importance, many aspects of papillomavirus biology are still obscure, and no specific anti-HPV drugs have been developed. Vaccines that inhibit infection by some HPV types are available, but vaccine access and uptake remain low in much of the world.

The papillomavirus particle is non-enveloped and consists of two viral proteins, L1 and L2, that form an icosahedral capsid containing viral DNA. The major capsid protein, L1, forms the shell of the capsid and binds to the cell surface, whereas the L2 protein mediates retrograde trafficking of the virus to the nucleus during virus entry. In each capsid, 360 molecules of L1 are assembled into 72 pentameric capsomers. 60 “hexavalent” capsomers each bind to six adjacent capsomers; the remaining “pentavalent” capsomers each bind five adjacent hexavalent capsomers. The structure of the L1 pentamer has been solved at high resolution by X-ray crystallography, and the structure of L1 in the capsid has been solved by cryo-electron microscopy (cryo-EM) (*1–4*). Pentavalent and hexavalent L1 pentamers both display five-fold symmetry with a pore extending entirely through the center of each pentamer from the outer to the inner surface of the L1 shell. The flexible C-terminal ends of each L1 monomer in the pentamer interact with adjacent pentamers to stabilize the capsid.

Each capsid also contains an estimated 12 to 72 L2 molecules (*5*). The structure of L2 in the capsid is not known. Early low resolution (∼22 Å) cryo-EM and biochemical analysis suggested that a single L2 molecule is present in each capsomer (*5, 6*), and a small density in the pore of pentavalent but not hexavalent bovine papillomavirus capsomers was speculated to represent L2 (*7*). However, L2 has not been identified in recent high resolution cryo-EM structures of the capsid, other than some small densities thought to represent short segments of L2 that have not been assigned to specific sequences of the protein (*2, 3, 8*). Antibody neutralization and binding studies suggest that much of L2 is accessible at the surface of various papillomaviruses (*9–11*).

Limited biophysical analysis of recombinant HPV16 L2 suggested that a significant amount of L2 is disordered (*12, 13*). AlphaFold2 predicts that most of the 473-amino acid HPV16 L2 protein is comprised of two intrinsically disordered segments, an ∼160-amino acid N-terminal disordered region (NDR, approximate extent, residues 68 to 224) and an ∼110-amino acid C-terminal disordered region (CDR, approximate extent, residues 333 to 446) (Fig. S1A, which shows the CDR and sequences to the end of the L2 protein as CDR+) (*14*). The functions of the disordered regions of L2 are not known, but the CDR acts in a sequence-independent manner to support virus entry (*14*). The NDR and CDR are separated by a central structured region (CSR, residues 224-333) that contains an anti-parallel β-sheet that binds cytoplasmic trafficking factors and chromatin during HPV entry (*14–16*). AlphaFold2 also predicts an α-helical segment near the N-terminus (up to residue 68). This overall L2 architecture has persisted for over 300 million years of papillomavirus evolution, implying these elements are functionally important (*14*).

After HPV binds to cells, L1 undergoes conformational changes that result in the exposure of the N-terminus of L2 for cleavage by furin protease (*17*). The virus particle is then internalized into the endosome, and a cationic cell-penetrating peptide (CPP) near the C-terminus of L2 mediates passage of this segment of L2 through the endosome membrane into the cytoplasm, where it binds cytoplasmic trafficking factors (*18*). Most of the L2 molecule is then thought to protrude into the cytoplasm until it is anchored in the endosomal membrane by an N-terminal transmembrane domain (*19, 20*). To insert into the endosome membrane, the C-terminus of L2 must emerge from the capsid. However, the molecular mechanism of L2 emergence and membrane passage is not known, although acidification of the endosome lumen and γ-secretase activity appear important for membrane insertion (*21–23*).

Here, we combine cryo-EM single particle analysis of purified HPV16 pseudovirus (PsV) particles with computational predictions to develop detailed models of the L2 protein structure and arrangement in the capsid. These models reveal heterogeneous related conformations of L2 in the capsid and suggest the route that the C-terminus of L2 takes to emerge from the capsid. This information may pave the way to anti-viral agents for HPV, which has so far defied drug development. Our results also provide new insights into how a well-structured protein can impose structure on intrinsically disordered protein segments, which may be broadly relevant.

## RESULTS

### Cryo-EM analysis of HPV16 capsids reveals the L2 protein associated with the inner surface of the L1 pentamer

To visualize L2 in the intact HPV16 capsid by cryo-EM, we studied density gradient-purified HPV16 PsVs, which consist of capsids composed of L1 and L2 and an encapsidated reporter plasmid and associated nucleosomes. We first studied infectious PsVs consisting of L1 and C-terminally FLAG-tagged L2 (L1/L2FLAG capsids) (*24*), as well as non-infectious PsVs that lack L2 (L1-only capsids). Examination of both PsV preparations by cryo-EM showed sheets of non-enveloped capsids with a diameter of ∼55 nm, the canonical size of HPV virions (Figure 1A,B). The capsids displayed icosahedral subunit structure, as expected, with minor heterogeneity in the shape of both L1/L2 and L1-only capsids, implying that HPV capsids have some inherent flexibility, consistent with published reports (*3, 8*).

**Figure 1.**
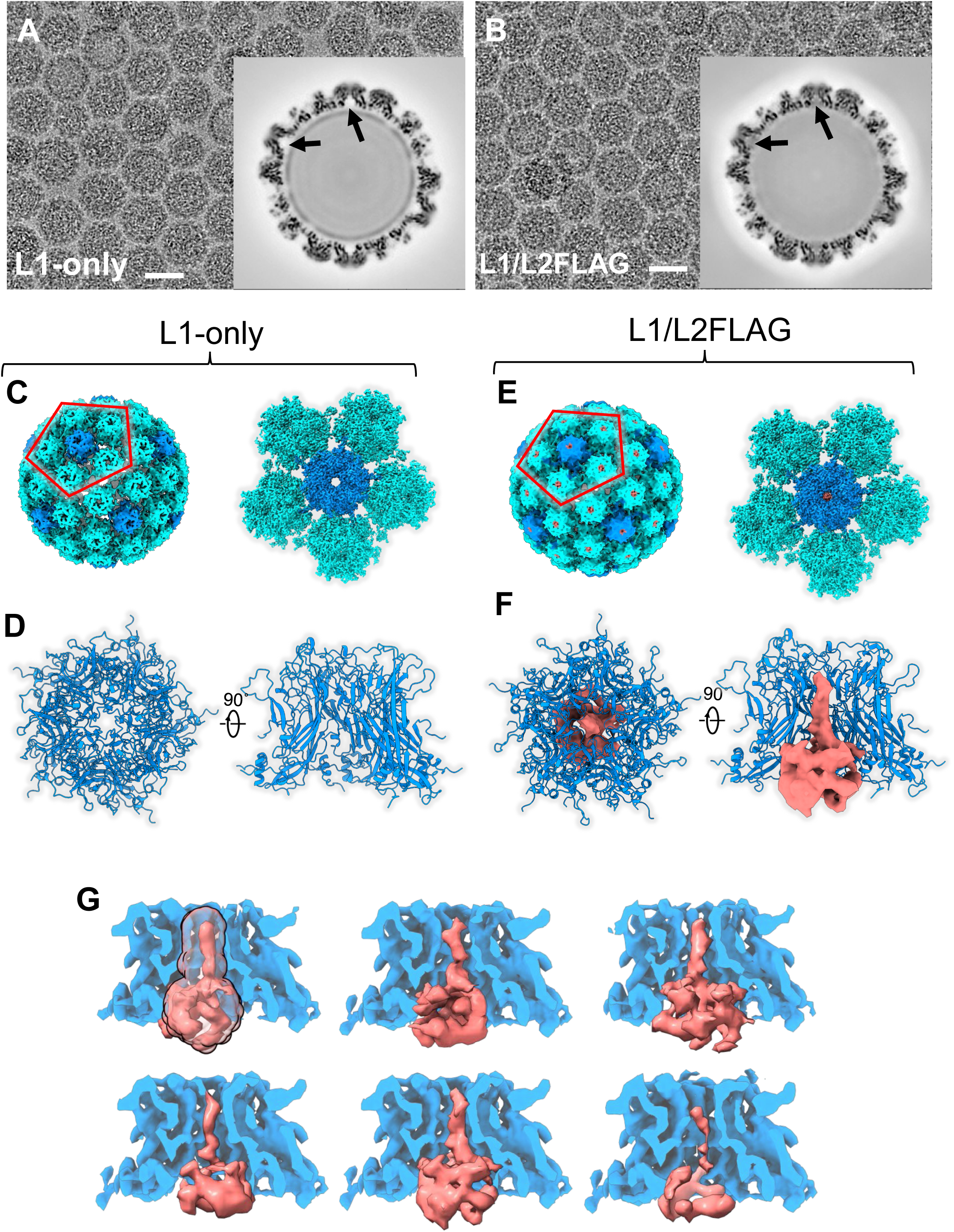
Cryo-EM reconstruction of HPV16 L1-only and L1/L2 capsids. A. Representative cryo-EM micrograph of the L1-only virus. The inset shows a central section from the 3D reconstruction. The arrows highlight the empty chamber formed by the L1 pentamer. Scale bar, 50 nm. B. Representative cryo-EM micrograph of the L1/L2FLAG virus. The inset as in panel A with the arrows highlighting the additional internal density within the pentamer chamber. C. 3D rendering of an L1-only capsid and a pentameric capsomer (dark blue) surrounded by five hexavalent capsomers (light blue). This color scheme is used in subsequent panels. The red pentagon outlines the five hexavalent capsomers surrounding a central pentavalent one. The capsomers were locally analyzed, revealing that the central pore of each capsomer is empty. D. Top view (left image) and side view with the front two L1 monomers removed (right panel) of pentavalent capsomers, shown in ribbon representation. E. 3D rendering of the L1/L2FLAG capsid as in panel C. Local analysis of capsomers revealed an additional internal density potentially corresponding to L2 protein, which is colored in pink in this and subsequent panels and figures. F. Top and side views of L1/L2FLAG pentavalent capsomers as in panel D. Potential L2 density represented in surface view by one of the selected 3D classes. G. Representative 3D classifications focused on the central densities within the pore and the L1 pentamer chamber in pentavalent capsomers in the L1/L2FLAG capsids. The transparent density in the upper left image indicates the mask region used for 3D classification. Side views are shown, with L1 in cross section.

To obtain high-resolution structures of HPV capsids, ∼3600 movie stacks for L1-only and L1/L2FLAG PsV samples were collected on a 300 kV electron microscope (Figure S1B,C). We generated initial 3D icosahedral structures of HPV capsids, and pentavalent and hexavalent capsomers were then extracted based on their orientations within the icosahedral symmetry and subjected to local structural reconstruction (Figure S1B,C; Table S1).

We solved the structures of the L1 pentavalent pentamer in the presence and absence of L2 to a resolution of 2.16 Å (PDB 37YB, EMD-78635) and 2.28 Å (PDB 37YL, EMD-78644), respectively. The corresponding hexavalent L1 pentamers were resolved at 2.37 Å (PDB 37YJ; EMD-78643) and 2.24 Å (PDB 37YM; EMD-78645). These structures are similar to previously published L1 pentamer structures (PDB: 7KZF) (*3*), with root mean square deviations (RMSDs) < 0.5 Å. Furthermore, RMSD of pentamers in the presence or absence of L2 is 0.124 Å (for pentavalent pentamers) and 0.161 Å (for hexavalent pentamers), indicating that the presence of the L2 protein has minimal effect on the structure of the L1 pentamer. As reported previously, L1-only pentamers contain an ∼43 Å-long open central pore with a minimum diameter of 18 Å that connects the exterior of the capsid with the dome-shaped space under the pentamer we designate the pentamer chamber (Figure 1A,C,D; Figure S2A). The flexible C-terminal arms of L1 that link adjacent capsomers were removed from the figures for clarity of presentation.

At low resolution, L1/L2FLAG capsids contain an amorphous density that is absent from L1-only capsids, which we attribute to the L2 protein (Figure 1A,B). This density is inside the pentamer chamber but is weaker than the overlying L1 pentamer in the preliminary 3D reconstructions of the L1/L2FLAG capsids, suggesting that it is more flexible than L1 and/or that it is present in only a fraction of capsomers. Further refinement revealed an ovoid L2 density that is approximately 32 Å high and 45 Å in diameter and fills the chamber of both pentavalent and hexavalent pentamers (Figure 1E,F; Figure S2B). Strikingly, a finger-like density extends on average ∼33 Å from the top of the L2 density in the pentamer chamber into the central pore of both pentavalent and hexavalent capsomers. The total density unique to L1/L2FLAG capsids is approximately 2/3 as large as expected for the tagged L2 protein (503 amino acids). All three-dimensional (3D) classes from both pentavalent and hexavalent L1/L2FLAG capsomers (see below) show this additional density, indicating that L2 is present in the great majority of capsomers. Similar results were obtained with PsVs containing an untagged version of L2 (PDB ID 37YN, EMD-78646 for pentavalent capsomers; PDB ID 37YO, EMD-78647 for hexavalent capsomers) (Figure S3).

Mass spectrometry of purified PsV preparations confirmed that L2 was present in only the L1/L2FLAG capsids and suggested that L1 was in approximately 6-fold molar excess over L2 (Figure S4), close to the expected 5-fold excess if one L2 molecule were associated with each L1 pentamer and consistent with the presence of the new cryo-EM density in most capsomers. Furthermore, other than histones and contaminating ferritin (which was abundant in both types of PsVs), other contaminants in the L1/L2FLAG capsid preparation were present in less than 15% the molar amount of L2 (Figure S4), which is not sufficient to account for the new density present in most capsomers.

The L2 density was resolved at approximately 8 Å (Figure 1F; Figure S2B), in contrast to the ∼2 Å resolution attained for L1 from the same set of virus particles. Hence, we could not assign specific amino acids of L2 to the cryo-EM density. Our inability to obtain a high-resolution structure of L2 suggests that the intrinsically disordered nature of L2 introduces substantial structural heterogeneity and/or conformational flexibility. Similarly, the N-terminal α-helices and the β-sheet identified by AlphaFold analysis of L2 alone were not resolved in the cryo-EM structure of L2 in the capsid, likely because they adopt variable orientations in space (see below).

Consistent with the absence of a unique dominant structure, focused 3D classification of capsomers from L1/L2FLAG capsids identified numerous distinct conformational classes that differ slightly in the overall morphology of the L2 density and the length and shape of the finger (examples shown in Figure 1G; Figure S2C; Figure S3E,G). No specific 3D classes dominated the population or were enriched in pentavalent compared to hexavalent capsomers, and rare classes lacked a finger (example shown in the lower right image of Figure S2C). In addition, there appear to be different densities that are discontinuous within the body of L2 in the various classes, possibly because different classes have different local segments with high structural heterogeneity that separate the densities that are visualized. As described in the next sections, AlphaFold3 modeling and molecular dynamics simulations generated models that align well with the cryo-EM density and provided additional evidence that L2 adopts heterogeneous structures in the capsid.

### AlphaFold3 models of HPV16 L2 associated with an L1 pentamer align well to the cryo-EM structures of the capsomer

To obtain additional evidence that the new cryo-EM density represents the L2 protein and to explore the molecular basis for the apparent structural heterogeneity of L2 in the capsid, we conducted AlphaFold3 modeling of one molecule of wild-type HPV16 L2 with five molecules of HPV16 L1 (*25*). AlphaFold3 predicted that L1 forms a pentamer that closely matches the L1 pentamer structure determined by cryo-EM (PDB:7KZF) (Figure S5). Because this analysis includes only a single capsomer, it does not distinguish between pentavalent and hexavalent pentamers, and the C-terminal tails of L1 dangle from the pentamer like tentacles from a jellyfish. The inclusion of L2 in the modeling had minimal effect on the predicted structure of the pentamer (RMSD <1 Å).

In the Alphafold3-predicted capsomers, L2 adopts a series of related low-confidence conformations. To assess the conformational diversity in these models, we used AlphaFold3 to generate 100 independent models of capsomers containing L2 associated with the L1 pentamer (*25*). The L2 protein in greater than 95% of these models aligns well with the cryo-EM density of L2 in capsomers (Figure 2A,B). We also conducted AlphaFold3 predictions of the L1 pentamer with ferritin and with several of the low abundance proteins contaminating the L1/L2 capsids identified by mass spectrometry. None of these proteins is predicted to align with the new cryo-EM density in the L1/L2FLAG capsomers (Figure S6).

**Figure 2.**
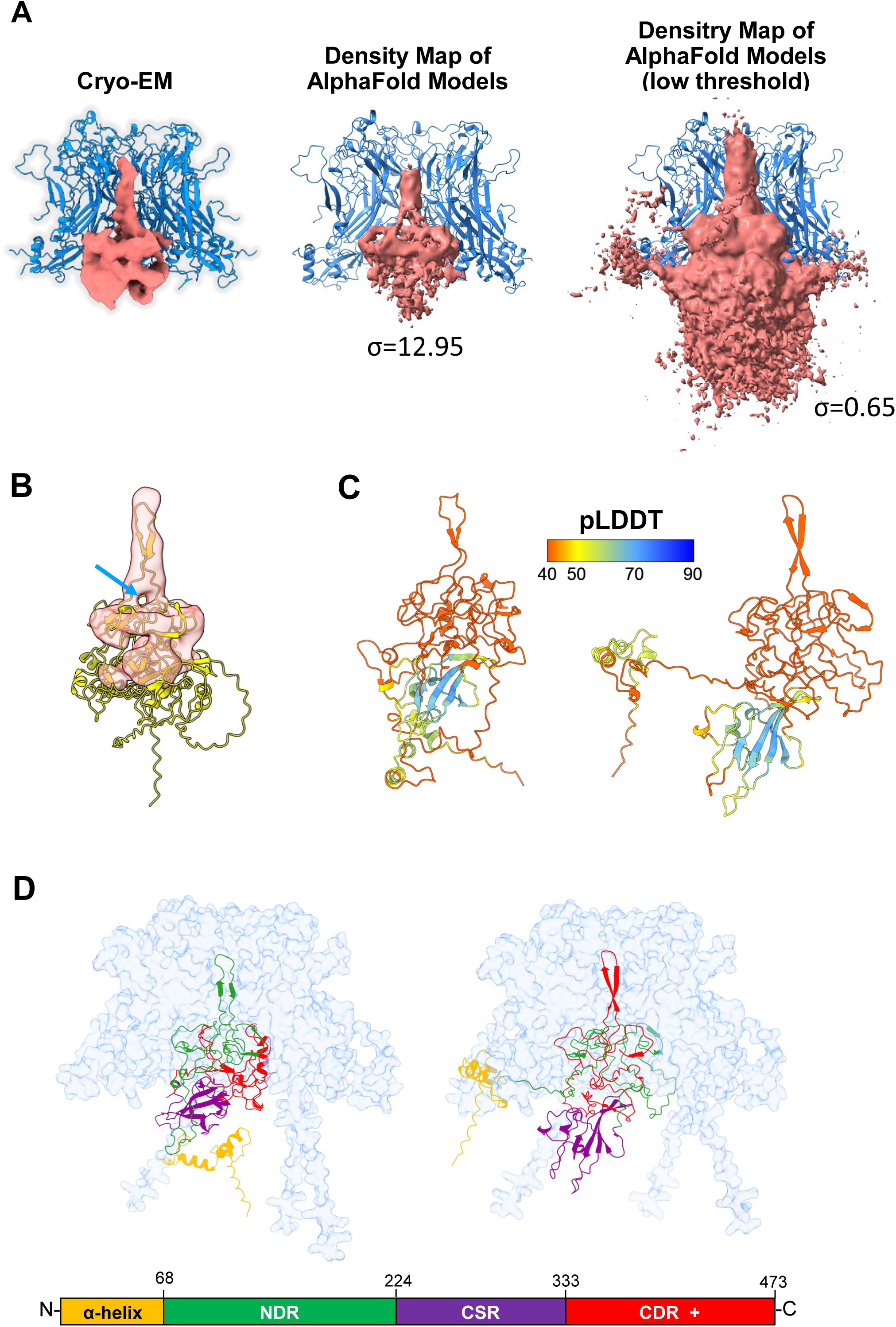
AlphaFold3 models of L2 in complex with L1 align well with the cryo-EM structure of L2 in L1/L2 capsids. A. Comparison of cryo-EM and Alphafold3 models. L2 structures are shown in pink, L1 pentamer is shown in blue ribbon view with the two L1 monomers in front of L2 hidden from view for clarity. Left image shows the same cryo-EM structure as in Figure 1G. The middle panel shows the 100 AlphaFold models of FLAG-tagged L2 converted into a density map and viewed at moderate threshold, which excludes highly variable parts of L2. The right image shows the AlphaFold density map viewed at a lower threshold, which emphasizes the heterogeneity of L2 structures beneath the pentamer chamber. Sigma value for the threshold level is indicated for each image. B. Superimposed cryo-EM density map and a representative AlphaFold3 L2 model. The pentamer used in the modeling is removed. The cryo-EM density of a 3D class is in translucent pink outline, and the AlphaFold3 prediction is colored yellow. The hole in the cryo-EM density at the base of the finger overlapping with the separation of strands in the predicted hairpin is highlighted with the arrow. C. Ribbon view of two AlphaFold3 models of HPV 16 L2 in the capsomer are shown in side view, with (right) or without (left) the N-terminal bulge. The pentamer used in the modeling is removed in these images. Colors represent pLDDT confidence scores, according to the color scale shown. Higher scores indicate higher confidence in the model. D. Ribbon view of two AlphaFold3 models of the pentamer-L2 complex are shown. Models with (right) or without (left) bulge are presented in side view. The pentamer (surface view) is colored light blue, and L2 (ribbon view) is color-coded by L2 domain as depicted at the bottom based on previous AlphaFold2 predictions of free L2 alone (*14*): N-terminal region (orange); N-terminal disordered region, NDR (green); central structured region, CSR (purple); and C-terminal disordered region including the C-terminus, CDR+ (red).

The L2 NDR and CDR, approximately half of L2, are predicted to form a compact mass that fills the pentamer chamber (Figure 2C,D). This mass aligns well with the cryo-EM density of L2, but the folded elements of L2 in the Alphafold3 predicted structures lie beneath the pentamer and their position is highly heterogeneous, most clearly visualized when a density map of the AlphaFold3 models is viewed at low threshold (Figure 2A, right image). Because of their heterogenous position, these folded segments of L2 are not visible in the cryo-EM structure and account for the “missing” L2 density.

Ninety-five of the 100 L2 AlphaFold3 predictions contain a finger-like projection from the disordered L2 domain extending into the pore of the pentamer, aligning with the finger in the cryo-EM density (Figure 2A-D). The fingers consist of sequences from the NDR or the CDR and most display a hairpin configuration consisting of amino acids extending from the top of the unstructured domain of L2, a short loop, and amino acids returning to the main body of L2 (Figure 2C-D). The separation of the leaving and entering strands of the hairpin in many AlphaFold3 models is consistent with the presence of a “hole” in the density at the base of the finger in some individual cryo-EM 3D classes (Figure 2B).

In six of 100 models, the finger consists of three strands of amino acids, a hairpin and a separate strand that ends at the extreme C-terminus of L2, pointing toward the outer surface of the pentamer (Figure 3A). In another four models, the finger ends at the C-terminus of L2, so the C-terminus is located within the pentamer pore or at its base. Finally, five models contained two hairpins in a single finger. Fewer than 5% of the models lack a finger and are not discussed further.

**Figure 3.**
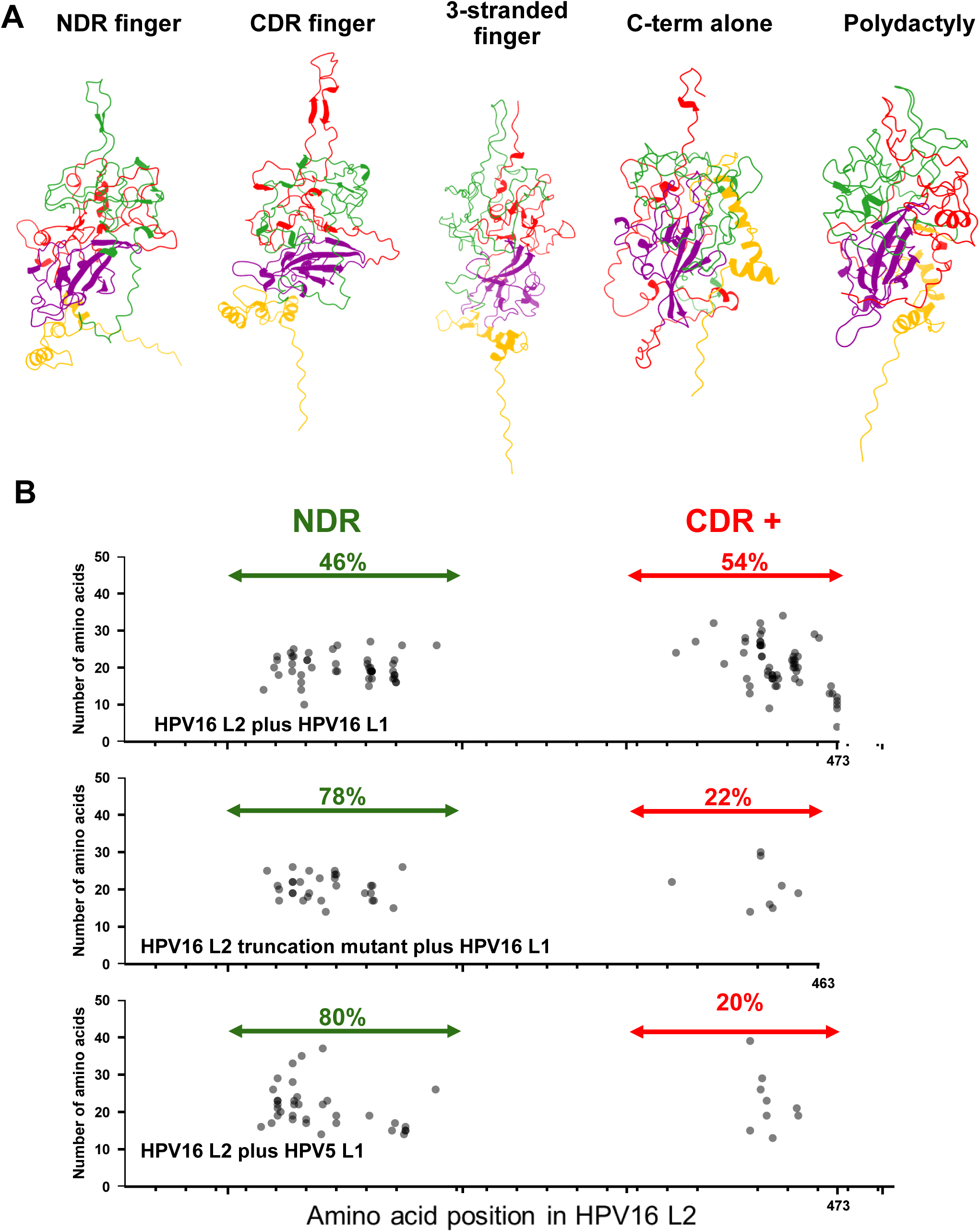
Diversity of finger configurations and positions. A. Five AlphaFold3 models of HPV16 L2 in the capsomer, illustrating variations in the finger. Each model is color-coded according to its position in L2 as in Fig. 2D. B. Position of fingers in HPV16 L2 AlphaFold3 models. The position of the “fingertip” in ∼100 models for the indicated combinations of L1 and L2 are plotted. Each dot shows the fingertip position and length of the finger (in number of amino acids) in an individual model. The NDR and CDR+ segments are shown by horizontal arrows, and the proportion of fingers in each segment is indicated. Models containing two hairpins or one hairpin and an additional strand in the finger were excluded from this analysis.

In the AlphaFold3 models, the N-terminal helical region of L2 is connected to the base of the unstructured domain of L2 in the pentamer chamber by the N-terminal portion of the NDR. In most models, the structure is underneath the pentamer, but in 25 of 100 models, this structure is at the periphery of the pentamer, forming a “bulge” that would presumably be positioned near an adjacent pentamer in the intact capsid (Figure 2C,D, right images). The β-sheet is predicted with high confidence (Figure 2C) and does not directly contact the pentamer and displays variable orientations in space as described below.

In AlphaFold3 models of the complex with either NDR or CDR fingers, the convex upper surface of the ovoid L2 mass is highly acidic due to the predominance of acidic amino acids in the NDR and CDR and primarily contacts basic residues in the concave inner surface of the pentamer chamber (Figure S7A,B). Examination of the predicted interface between L1 and L2 showed that amino acids throughout the NDR and CDR of L2 interact with L1, with few interactions involving the CSR or the N-terminal helices (Figure S7C).

Similar AlphaFold3 models were obtained for three other divergent papillomaviruses: HPV5, a cutaneous HPV type (unlike HPV16, which infects mucosa); bovine papillomavirus type 1, a distantly related papillomavirus that causes skin fibropapillomas; and CmPv1, the most divergent known papillomavirus, which infects sea turtles (Figure S8). This finding suggests that the main structural features of L2 in the capsid are conserved among all papillomaviruses.

### AlphaFold3 predicts marked variability of the position of the L2 finger, the structure of the disordered regions, and the orientation of the folded elements of L2

In examining many independent models of untagged L2 in capsomers, AlphaFold3 predicted that the positions of the fingers along L2 are distributed throughout the disordered segments of the protein, with approximately equal representation of fingers in the NDR and in the CDR to the C-terminus of the protein (a segment designated CDR+) (Figure 3B). No finger contained sequences from the N-terminal helices or CSR. The length of the fingers also varied in different models from 4 to 34 amino acids, but none extended through the pore to the external surface of the pentamer. The L2 proteins from diverse papillomavirus are also predicted to contain fingers at various positions in the NDR and CDR when associated with the cognate L1 pentamer, although the relative proportions of NDR and CDR fingers varied among the different virus types, with HPV5 being the most biased, with 95% of fingers arising from the NDR (Figure S9).

To determine whether the sequence of L2 affects the predicted location of fingers, we modeled an infectious HPV16 L2 mutant that lacks the C-terminal 10 amino acids of L2. As shown in Figure 3B middle panel, 78% of fingers in the mutant are located in the NDR, instead of being distributed approximately equally between the NDR and the CDR as is the case for wild-type L2. We also modeled HPV16 L2 associated with HPV5 L1 pentamers. As shown in Figure 3B bottom panel, instead of the equal distribution of fingers between the NDR and the CDR+ when wild-type HPV16 L2 associated with the cognate HPV16 L1 protein, 80% of the fingers were from the NDR when HPV16 L2 was modeled with HPV5 L1. Thus, the sequence of both L1 and L2 influences the position of the fingers in L2 in the AlphaFold3 predictions of the L1/L2 capsomer.

We also used AlphaFold3 to predict the variability of different segments of L2 in models of the HPV16 L1/L2 complex. For this analysis, we separately aligned the N-terminal helical domain, the NDR, the CSR, and the CDR+. As shown in Figures S10A and B, the structures of both the NDR and the CDR are highly variable (RMSD > 18 Å in pairwise comparisons), even when the fingers are at similar positions along the length of L2. In contrast, the folded structures of the CSR and the N-terminal helices are very similar in different models (RMSD typically < 2 Å), but these elements swivel and tilt relative to the positions of the fingers (Figure S10C).

### Association of L2 with the L1 pentamer stabilizes L2

We also conducted all-atom molecular dynamics (MD) simulations of a representative AlphaFold3 model of an HPV16 capsomer with L2 containing a finger with a tip at position 443 in the CDR. Two replicates of this model were simulated for at least 500 ns, with similar results. The main structural features of the complex, namely the presence of a finger and the NDR and CDR filling the pentamer chamber, persisted over 500 ns simulation, indicating that the predicted structure is plausible and stable over this timeframe (Figure 4A). L2 is more dynamic in the complex than L1, consistent with its disordered nature (Figure 4B, green and blue lines). The N-terminal helical domain and β-sheet remain folded but have high root mean square fluctuation (RMSF) values because they are beneath the pentamer chamber and adopt different positions in space, whereas the disordered regions of L2 are less flexible because they are constrained inside the pentamer chamber and pore (Figures 4A,C). We also conducted MD simulations of L2 in the absence of L1. Free L2 was far more flexible and less densely packed than L2 associated with the pentamer (Figures 4B, red lines,D-G). The increased dynamics of free L2 compared to L2 in complex with L1 was more dramatic for the NDR and CDR than for the CSR (Figure 4F).

**Figure 4.**
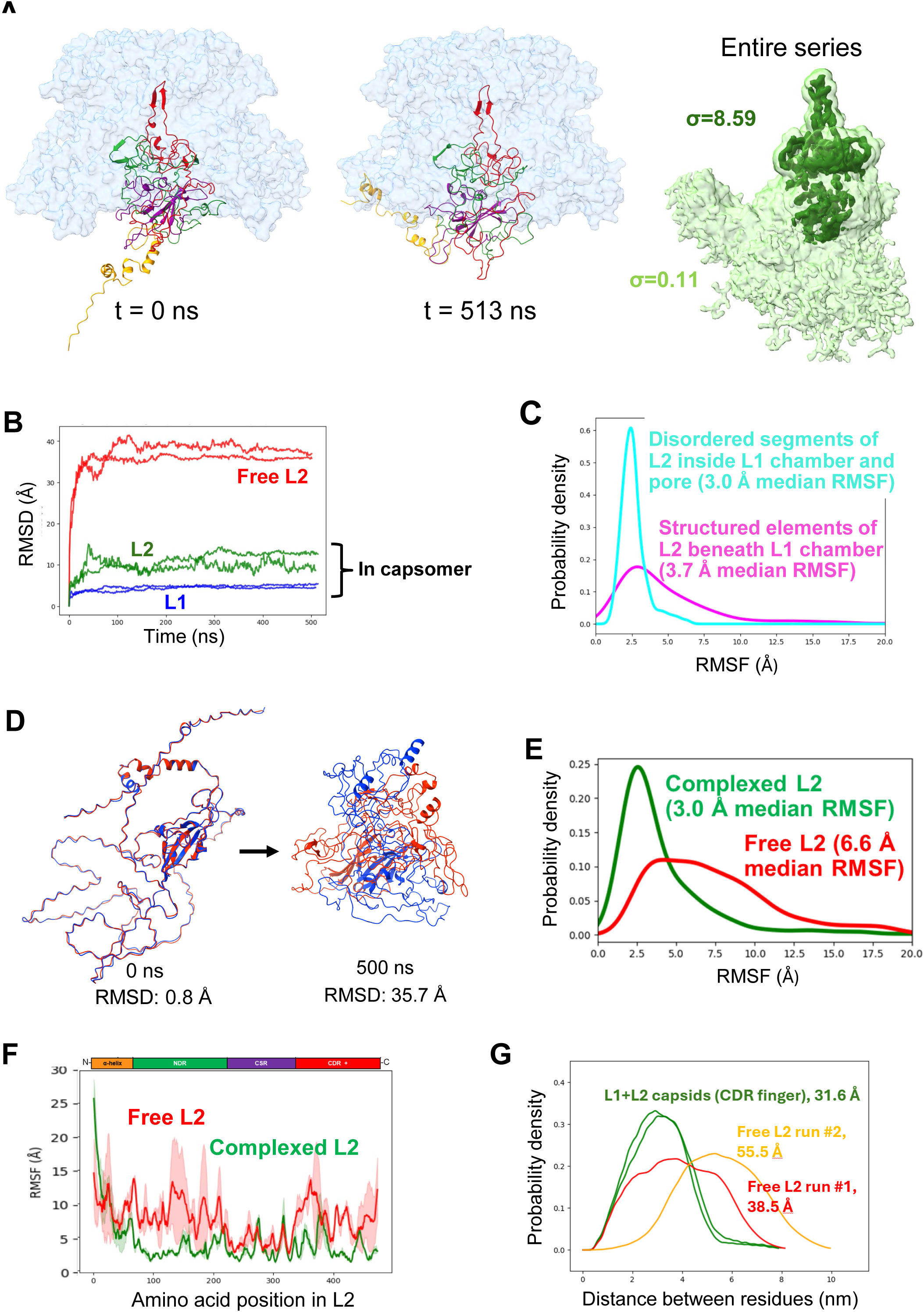
Molecular Dynamics simulations reveal that L2 is more compact and less flexible when associated with L1 pentamer. A. Left and middle images show snapshots of an all-atom molecular dynamics simulation conducted for an AlphaFold3 model of L2 with a fingertip at position 443 in the CDR associated with a pentamer consisting of truncated L1 (positions 1 to 480 for L1, lacking the 25 C-terminal amino acids). The ribbon diagrams show a side view of the starting structure of L2 on the left and after 513 ns simulation in the middle, color-coded as in Figure 2D. L1 is shown in light blue surface representation. Image on right shows superimposed L2 density maps of simulation time points sampled every 10 ns for each of two replicate 500 ns simulations. Threshold values for each image, shown in light and dark green, are noted as the sigma value. B. RMSD time series of L1 (blue) and L2 (green) compared to the starting frame in two >500 ns replicate simulations of the complex containing L2 with a finger in the CDR, showing that the simulations reach equilibrium and that L2 changes more from its starting structure than L1. The lines in red show a similar time series of simulations of L2 in the absence of L1. C. Estimated probability density function of RMSFs for the disordered segments of L2 inside the L1 pentamer chamber or pore (teal) or the structured elements of L2 beneath the pentamer chamber (lavender). Estimated based on RMSF distributions from replicate ≥ 500 ns simulations of L2 containing a CDR finger, using a Gaussian kernel density estimation in SciPy. D. Snapshots of two replicate all-atom molecular dynamics simulations (red and blue) from the same starting AlphaFold3 structure of L2 in the absence of L1 at 0 ns and 500 ns, with the average RMSD between the simulations shown for both timepoints. E. Estimated probability function showing RMSF of free L2 (red) or L2 associated with an L1 pentamer (green), calculated from the simulations shown in panels A and B, respectively. Because of the rapid initial reorganization of free L2 (see panel B), RMSF calculation for free L2 excluded the first 50 ns. F. RMSF of L2 in simulations of L2 alone (red) and in complex with the L1 pentamer (green), according to the amino acid position in L2. The solid line is the average RMSF of two replicate simulations and the shaded area is the standard deviation. G. Estimated probability function of pairwise distances between residues in the NDR and in the CDR, sampled every 0.5 ns from 200 to 500 ns in replicate MD simulations of a model of the L1/L2 complex with a finger in the CDR (green) and a model of free L2 (red and yellow). The values for the L1 + L2 capsids are the averages of the two replicates for each model. Because of the variability in the results for the model of free L2, the value for each replicate is shown separately.

To assess compactness of the NDR and CDR during MD simulations, we measured the pairwise distances between each backbone Cα carbon atom in the NDR and each Cα atom in the CDR. We compared these distances in models of the complex with a finger in the CDR to models of free L2. This analysis (Figure 4G) showed that in the absence of L1, amino acids in the NDR of L2 were on average 3.85 and 5.55 nm away from the amino acids in the CDR in replicate simulations, but in the presence of L1, this distance was reduced to ∼3 nm, with little variability in replicate simulations. Thus, the disordered domains in L2 are closer together when complexed with the L1 pentamer than in the absence of L1. Overall, these results indicate that the association of L2 with the pentamer forces the disordered segments of L2 to adopt relatively stable, compact configurations inside the pentamer chamber and pore.

### The C-terminus of L2 is exposed on the surface of the capsid

We next conducted AlphaFold3 modeling of the FLAG-tagged HPV16 L2 variant used for the cryo-EM studies shown in Figure 1. AlphaFold3 predicts that FLAG-tagged L2 in complex with the pentamer retained the major structural features described above for untagged L2, with approximately 10% of fingers containing the C-terminal tag itself (Figure 5A). Notably, in six of 100 models the C-terminal FLAG segment extends entirely through the central pore of the pentamer and emerges from the outer surface of the pentamer (Figure 5B, right panel). In two additional models, the N-terminus of L2 emerges through the pore (Figure 5B, left panel).

**Figure 5.**
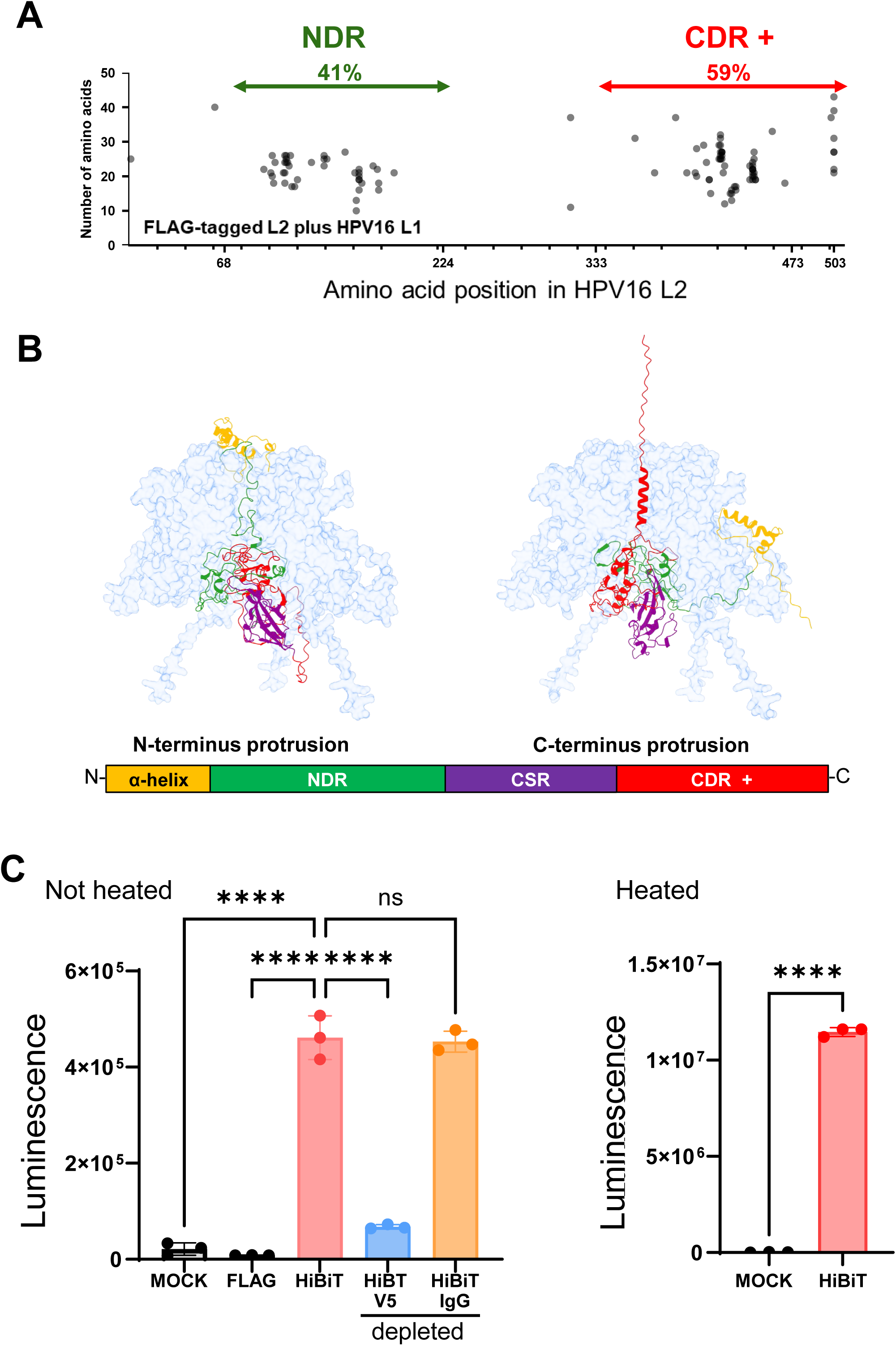
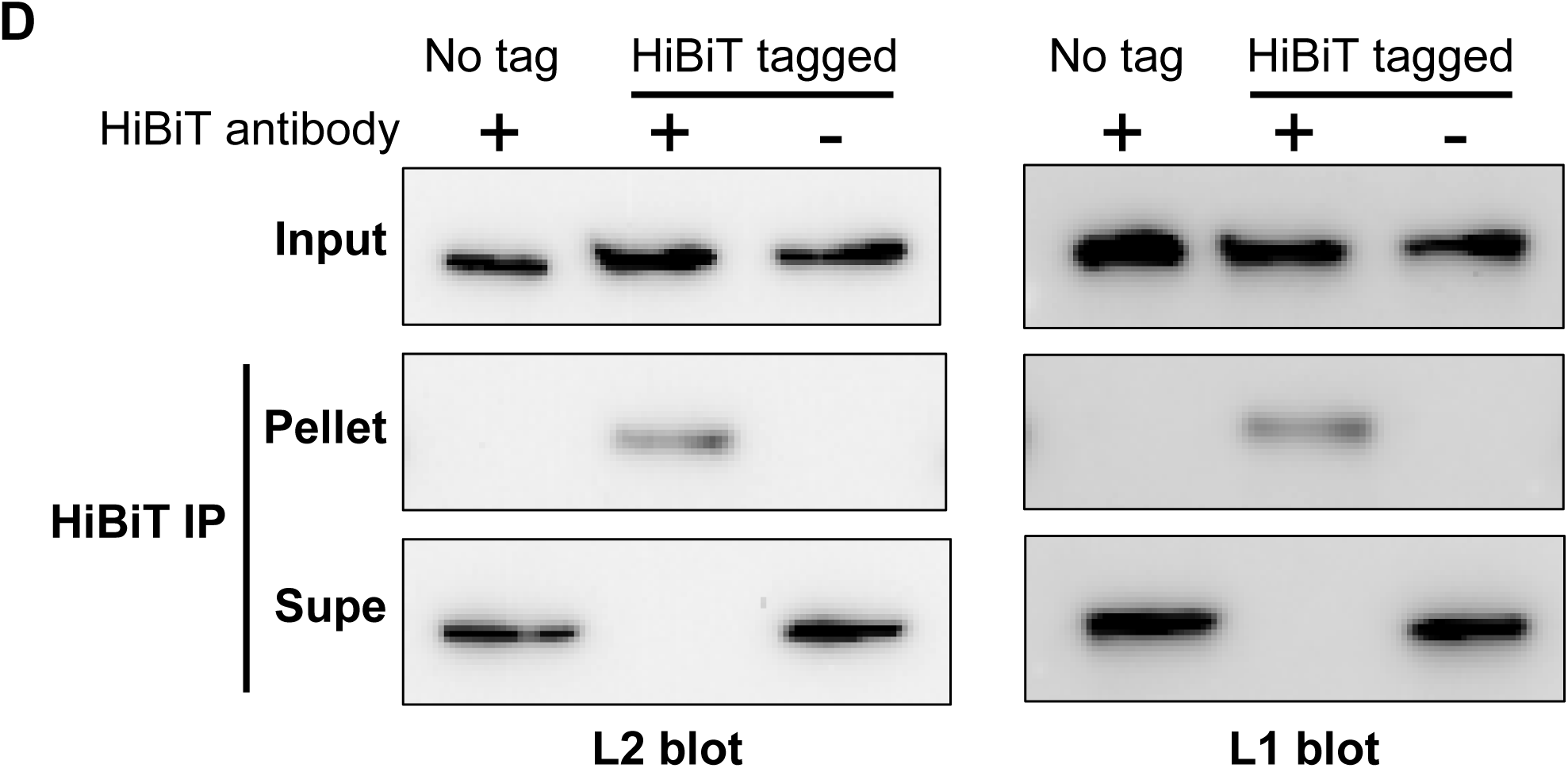
C-terminus of L2 is exposed on the surface of the capsid. A. Distribution of fingers of FLAG-tagged HPV16 L2. Location of finger in AlphaFold3 models of HPV16 L1 and HPV16L2 with a C-terminal 3x FLAG tag. The position of the fingertip from multiple AlphaFold3 predictions are plotted as in Figure 3B. B. Emergence of L2 from the pentamer pore. AlphaFold3 predictions were performed for C-terminally FLAG-tagged L2 with wild-type HPV16 L1 pentamer. Representative models with emergence of the N-terminal helices (left) or C-terminal segment (right) are shown. L1 and L2 colored as shown in Fig. 2D C. Split luciferase assays on HiBiT-tagged PsVs. Left panel. Purified HPV16 PsVs containing either a FLAG tag or a HiBiT tag at the C-terminus of L2 were incubated with anti-L1 monoclonal antibody (V5) or a negative control mouse antibody (IgG). PsV– antibody complexes were pulled down using Protein G-coated magnetic beads and the resulting supernatant was assayed for luminescence activity in the presence of LgBiT, expressed as relative light units. Mock refers to no PsV added. Right panel. Purified HPV16 PsVs containing a HiBiT tag at the C-terminus of L2 was assayed as in left panel except antibody binding was not performed and the samples were heated prior to assay to assess the total amount of L2 present in the sample. Statistical analysis was performed using two-way ANOVA in Prism version 11 (GraphPad Software). Statistical significance was defined as follows: ns, not significant; ****, p < 0.0001. D. Co-immunoprecipitation of capsids with antibody recognizing the C-terminal HiBiT tag on L2. Purified HPV16 PsV containing either untagged L2 or L2 with a C-terminal HiBiT-tag were incubated with or without anti-HiBiT antibody. Immune complexes were captured using Protein G-coated magnetic beads. Input, supernatant (Supe), and eluate (Pellet) fractions were analyzed by western blot for levels of L1 (right panel) and L2 protein (left panel). Data are representative of three independent experiments.

These results imply that the C-terminus of the L2 protein can emerge through the pore. To test if the C-terminus of L2 was accessible on the exterior surface of intact capsids, we first used a split luciferase assay as a measure of exposure of the C-terminus of L2 from the capsid. In this assay, luciferase activity is reconstituted when a short inactive fragment of luciferase, named HiBiT, associates with a long inactive fragment named LgBiT. We appended HiBiT to the C-terminus of the L2 protein and produced infectious PsVs containing this L2 variant. We then mixed gradient-purified HiBiT-tagged PsVs with extracts of cells expressing LgBiT and used reconstitution of luciferase activity as a measure of exposure of the C-terminus of L2 on the exterior of the capsid. As shown in Figure 5C left panel and S11A, addition of HiBiT-tagged PsV to extracts containing LgBiT caused a dose-dependent increase in luciferase activity, whereas PsVs containing a C-terminal FLAG tag instead of HiBiT did not induce activity. Depletion of intact capsids from the PsV preparations by immunoprecipitation with anti-L1 antibody V5 eliminated luciferase activity, but immunoprecipitation with control IgG did not, indicating that the activity induced by the PsV preparation was due to L2 associated with L1 (Figure 5C, left panel). Furthermore, heating the samples prior to luciferase assay to release L2 from inside the capsid caused an approximately 25-fold increase in luciferase activity (Figure 5C, right panel), suggesting that the C-terminus of most L2 molecules in intact capsids is unavailable to associate with LgBiT and reconstitute luciferase activity.

To determine whether the majority of capsids contained L2 exposed on their surface, we used an anti-HiBiT antibody for co-immunoprecipitation experiments with purified PsV containing L2 with a C-terminal HiBiT tag. As shown in Figure 5D, anti-HiBiT immunoprecipitated all of L2 and L1 from PsV preparations containing HiBiT-tagged L2. In contrast, L1 was not immunoprecipitated from capsids containing untagged L2. This result implies that the C-terminus of at least one L2 molecule is exposed on the surface of essentially all capsids.

### The C-terminus of multiple L2 molecules is preferentially exposed on the surface of each capsid

Finally, to determine if the C-terminus of multiple L2 molecules emerged from morphologically intact capsids, we conducted immunogold staining and transmission electron microscopy (TEM). We prepared infectious PsVs containing L2 lacking an HA tag or with an HA tag at its C-terminus. PsV were adsorbed to EM grids, stained with anti-HA antibody followed by secondary antibody coupled to gold nanoparticles, and imaged by TEM after negative staining. In the experiments shown in Figures 6A and B, numerous intact, well-formed icosahedral capsids were observed. Virtually all capsids containing C-terminally-tagged L2 were stained, and on average more than seven electron-dense gold particles were observed at or within a few nanometers of each capsid. In contrast, untagged capsids showed minimal staining (∼0.2 gold particle/capsid), as did capsids with C-terminally tagged L2 if the primary antibody was omitted.

**Figure 6.**
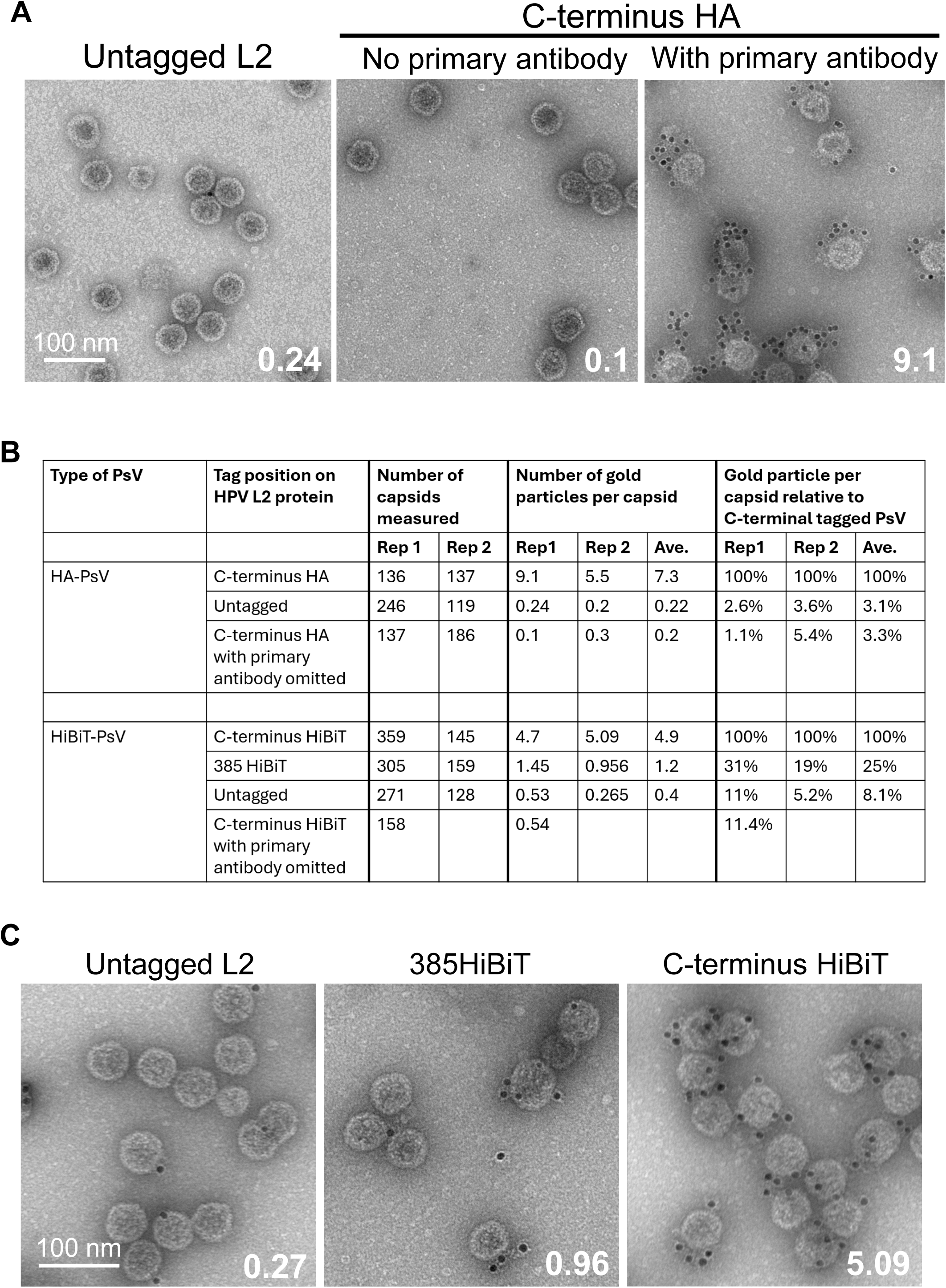
Transmission electron microscopy visualizes C-terminus of L2 on the surface of intact capsids. A. Immunogold transmission electron microscopy of HPV16 PsV particles. Transmission electron micrographs of purified HPV16 PsVs containing an HA tag at the C-terminus of the L2 protein or untagged L2. PsV were labelled with anti-HA primary antibody and a gold-conjugated goat-anti-mouse secondary antibody. The anti-HA primary antibody was omitted where noted. Samples were negatively stained with uranyl formate and imaged. The numbers indicate the mean number of electron-dense gold particles observed per capsid. B. Table summarizing immuno-gold staining of pseudovirus. Rep 1 and Rep 2 refer to independent biological replicates. Percent gold tags per capsid are normalized to the number of particles observed for C-terminally tagged L2, set at 100%. Ave. shows the average number of gold particles per capsid and the average percentage of gold particles per capsid normalized to capsids containing the C-terminal tag. C. Transmission electron microscopy of pseudoviruses stained with anti-HiBiT antibody. PsV preparations with no HiBiT tag or a HiBiT tag at the L2 C-terminus or position 385 were stained with anti-HiBiT antibody followed by gold-labeled secondary antibody, as in Fig. 6A. The anti-HiBiT primary antibody was omitted where indicated.

We also examined L2 exposure on PsV containing a HiBiT tag at L2 position 385 in the CDR or at the C-terminus. Both of these viruses are infectious (Figure S11B), and the anti-HiBiT antibody bound the tag at position 385 at least as well as the tag at the C-terminus as assessed by western blotting (Figure S11C). Intact capsids containing a C-terminal HiBiT tag on L2 showed on average 5 gold particles per capsid when stained with an anti-HiBiT antibody, whereas capsids with the HiBiT tag at position 385 showed only 25% as many gold particles, with many capsids remaining unstained (Figures 6B,C, S11D). These results show that morphologically intact capsids contain multiple copies of the L2 protein with a C-terminus accessible to antibody and suggest that a portion of the CDR upstream of the C-terminus was exposed in fewer capsids.

To provide addition evidence that the C-terminus of L2 was preferentially exposed compared to upstream segments of the protein, we also performed antibody neutralization experiments with PsV described above lacking a HiBiT tag or containing the tag at the C-terminus of L2 or at position 385. To measure the effect of the antibody on infection, we incubated anti-HiBiT antibody with PsV, added the PsV to HeLa cells, and three days later measured HcRed fluorescence expressed from the packaged reporter plasmid as a measure of infectivity. As shown in Figure S11B, left panel, the HiBiT antibody neutralized infection by the PsV containing the HiBiT tag at the C-terminus of L2. In contrast, PsV lacking the tag or containing the HiBiT tag at position 385 were not neutralized. Similarly, anti-HA antibody neutralized infection by PsV containing a C-terminal HA tag (Figure S11B, right panel). Taken together, these results suggest that the C-terminus of L2 is exposed on the surface of the capsid, whereas position 385 is exposed in few if any capsids.

## DISCUSSION

An understanding of the structure of any papillomavirus L2 capsid protein has long been elusive. Here, we use a variety of approaches to visualize L2 in infectious HPV16 PsV capsids. All or most capsomers in HPV16 PsV contain a single L2 molecule, but the occupancy of authentic HPV16 has not been assessed. The presence of L2 in each capsomer is consistent with prior low resolution (22.7 Å) cryo-EM analysis of HPV16 PsV (*5*). Our results show that L2 does not adopt a single, fixed conformation within the capsid. Rather, due to its intrinsically disordered nature, L2 exists as an ensemble of conformations with a similar overall shape. The disordered regions of L2 are constrained within the pentamer chamber and form a finger-like projection extending into the central pore of the pentamer such that the C-terminus of multiple L2 molecules are poised to emerge from the capsid through the pore and insert into the endosome membrane. The intrinsic disorder of L2 prevents the assignment of specific L2 amino acids sequences to the cryo-EM density. Nevertheless, the multiple related conformations of L2 predicted by AlphaFold3 align well with the cryo-EM density, leading us to conclude that these predictions accurately reflect the conformations of L2 in the capsid.

AlphaFold3 modeling and MD simulations indicate that in the absence of L1, L2 is largely disordered other than the N-terminal α-helices and the central β-sheet. When L2 is bound to the L1 pentamer, it adopts multiple conformations whose overall shape is similar in the vast majority of models. The disordered segments of L2 inside the pentamer chamber and pore are more compact and less flexible than they are in free L2. The upper disordered surface of L2 adopts a convex shape that binds the inner surface of the pentamer, and a finger extends from the main body of L2 into the central pore of the pentamer. The best-folded segments of L2, the α-helices and the β-sheet, are predicted to remain folded but adopt many different orientations distal to the inner surface of the pentamer, explaining our inability to resolve these elements by cryo-EM.

Despite the overall shared shape of multiple L2 structures predicted by AlphaFold3 and inferred from the cryo-EM results, the configurations of the disordered regions vary widely in the L2 models. Most strikingly, many different L2 sequences throughout the disordered segments can form fingers in the pore. The position of the fingers is determined in part by the sequence of L1. Interestingly, the disordered segments of L2, which are the most flexible segments in free L2, are less flexible than the structured elements when complexed to L1. These unusual features suggest that the L1 pentamer largely determines the structure of the bound L2 protein: the flexible L2 sequences conform to a mold formed by the pentamer chamber and pore. More generally, our results show that a rigid scaffold, in this case the L1 pentamer, can impose a relatively stable structure on long disorder protein segments.

Our results provide insights into the action of L2 during virus entry. Both ends of L2 are thought to emerge from the capsid early during virus entry even though the capsid appears to remain relatively intact during this process (*26*), implying that L2 emergence does not cause major structural changes in the capsid. The emergence of the N-terminal helices through the central pentamer pore was observed in rare models of L2 with a C-terminal FLAG tag, suggesting that the N-terminus of L2 may be able to emerge from the pore. However, the location of the N-terminal helical segment at the periphery of the pentamer in a larger fraction of models suggests that it might be also able to emerge between pentamers, possibly after reduction or isomerization of disulfide bonds that link neighboring pentamers, cleavage of L1 by cellular proteases, or heparin binding, which causes the L1 arms that link adjacent capsomers to become more extended (*27, 28*).

Importantly, our results provide a route for the C-terminus of L2 to emerge from the capsid, a critical step required for L2 to penetrate the endosome membrane and initiate retrograde trafficking of the incoming virus particle during HPV entry. In AlphaFold3 modeling, we observed the C-terminus of L2 in the pentamer pore – either free-standing or as part of a three-stranded structure (Figure 3A) – or actually emerging through the pore to the outer surface of the pentamer (Figure 5B). Consistent with this model, a series of experiments with purified PsV showed that multiple copies of the C-terminus of the L2 protein are exposed on the surface of the great majority of capsids, whereas more upstream segments of L2 are less often exposed.

Because the disordered nature of L2 has persisted over papillomavirus evolution, this arrangement must be advantageous to the virus, but why does L2 employ multiple conformations instead of utilizing a fixed structure with its C-terminus in the pore of every capsomer? We recently found that the presence of a structured element in the CDR inhibits protrusion of L2 through the endosome membrane (C.O. and D.D., manuscript in preparation). This finding suggests that a folded structure that dictates a fixed conformation of L2 with its C-terminus in the pore is incompatible with membrane protrusion. Hence, we hypothesize that L2 evolved to have unstructured disordered segments to allow membrane protrusion, thus necessitating a fluid approach to find a productive conformation that allows emergence from the capsid. Alternatively, it may be deleterious to have the C-terminus of too many L2 molecules emerge from the capsid, or the existence of multiple L2 configurations within the capsid may reflect selection for conformational flexibility that enables L2 to execute its roles during virus entry after emergence from the capsid, including the ability to bind multiple cellular trafficking factors.

We imagine two basic ways L2 may adopt a productive configuration. Each L2 molecule may transiently sample a variety of fingers until a conformation conducive to externalization of L2 is attained. Alternatively, each L2 molecule in the capsid may have a unique conformation, but because the capsid contains many L2 molecules, it is likely there will be one or more in a productive conformation in every capsid. In either scenario, the existence of multiple molecules of L2 in each capsid enables many L2 configurations to be sampled by the virus.

Most of L2 is not visible in recent published cryo-EM structures of the HPV16 capsid (*1–3*). It is possible that L2 exists in two major configurations, analogous to pre-fusion and post-fusion forms of enveloped virus fusion proteins. One configuration is largely inside the capsid as reported here and the other is invisible to cryo-EM analysis because it is unstructured after it has emerged from the capsid. The existence of L2 in either a pre-or a post-triggered form reconciles the structure reported here with indirect evidence that N-terminal segments of L2 are exposed at the surface of the capsid. The nature of the trigger remains to be determined, but our models provide a possible mechanism. Unlike the roof of the pentamer chamber, which is basic and contacts the acidic disordered mass of L2, the outer segment of the pentamer pore is highly acidic (Figure S7B). This acidic segment may impede the passage of the highly basic C-terminal CPP through the pore. Acidification of the endosome lumen following internalization of the virus is likely to protonate these residues in the pore and facilitate emergence of L2 from the capsid.

The structure of L2 in the capsid suggests possible new anti-viral approaches. Because the C-terminus of L2 exits through the central pore of the pentamer, peptides or small molecules that constrict or occlude the pentamer pore might block L2 emergence from the capsid and inhibit infection. Similarly, approaches that stabilize intercapsomeric interactions may prevent emergence of the N-terminal helices. The heterogenous, dynamic nature of L2 suggests alternative strategies to inhibit L2 emergence, namely to stabilize non-productive conformations of L2 or restrict the flexibility of L2. Finally, understanding the structure of L2 may facilitate its use as a broadly protective HPV vaccine (*29*).

## Methods and Materials

### Cells and cell cultures

HeLa S3 cervical carcinoma cells were purchased from American Type Culture Collection (ATCC). HEK293TT cells, which were derived from HEK293T cells by overexpressing SV40 large T antigen, were obtained from Christopher Buck (National Institutes of Health). All cells were grown with Dulbecco’s modified Eagle’s medium (DMEM, Gibco) with HEPES and L-glutamine, supplemented with 10% fetal bovine serum (FBS) and 100 units/mL penicillin streptomycin, at 37°C in 5% CO_2_. Cell lines were authenticated by the ATCC cell authentication service.

### Plasmids and PsV production

Packaging plasmids designated p16sheLL, expressing HPV16 L1 only or L1 plus L2 (either untagged or containing an epitope tag) were used to produce wild-type PsV (*30*). The PsV containing a C-terminal 3xFLAG tag was described previously (*24*). The C-terminal HA tag in L2 was inserted downstream of a C-terminal 3xFLAG tag and a cleavage site for thrombin and was described previously (*24*). A HiBiT tag [VSGWRLFKKIS] was inserted in-frame at L2 position 385 or a GGSGGGSG linker following by the HiBiT tag was inserted at the C-terminus of L2.

5 x 10^6^ 293TT cells were seeded per plate in 15 cm dishes and incubated at 37°C. After 24 h, polyethyleneimine was used to co-transfect cells with a reporter plasmid [pCAG-HcRed (Addgene #11152)] and p16sheLL expressing HPV16 L1 alone (to generate L1-only PsV) or L1 and L2. After 72 h, the cells were harvested, mixed with (phosphate-buffered saline [PBS] containing 0.5% Triton X-100, 10 mM MgCl₂, 5 mM CaCl₂, and 0.1% RNase A) and incubated overnight at 37°C. Packaged PsVs in the cell lysate were purified by density gradient centrifugation in Optiprep, and gradient fractions containing the most L1 as assessed by SDS-polyacrylamide gel electrophoresis and Coomassie brilliant blue staining were pooled and stored at –80°C (*30*).

### Cryo-EM sample preparation and data collection

The HPV16 L1/L2FLAG and L1-only PsV samples were assessed for purity and concentration before vitrification for cryo-EM data collection on the Titan Krios. 3 µl of the purified virus sample (approximately 3 μg PsV) was pipetted onto a glow-discharged Quantifoil Cu R2/1 300 mesh grid, blotted for 5 s under 90% humidity and rapidly plunge-frozen in liquid ethane using a Leica EM GP2 automated freezer. Movie stacks were collected on a Titan Krios G2 microscope (Thermo Fisher Scientific, Waltham, MA) equipped with a field emission gun, a K3 Summit direct-detection device (Gatan, Pleasanton, CA), and an energy filter with a slit width of 20 eV. Automated data collection was performed using SerialEM (*31*) with a typical defocus range of –1.6 to –2.2 μm and total dose of 55 e^−^/Å^2^ per movie, with a total exposure time of 2 s, yielding 40 frames (0.07 s/frame) per image stack. In total, 3,660 and 3,594 movie stacks were collected for L1/L2FLAG and L1-only PsV, respectively, at a nominal magnification of ×81,000 (calibrated pixel size: 0.534 Å).

### Cryo-EM data processing

Preprocessing was performed using cryoSPARC v4.7.0 and v4.7.1 (*32*). Movie stacks of L1/L2FLAG PsV were motion-corrected using the Patch Motion Correction tool with a 1/2 crop factor, resulting in a binned pixel size of 1.068 Å. Contrast transfer functions (CTFs) were estimated using Patch CTF Estimation. A manually curated subset of 326 capsids was used for 2D classification to generate templates for Topaz training (*33*).

This enabled automated particle picking via Topaz Extract, yielding an initial set of 228,327 high-quality capsids. The motion-corrected micrographs and selected capsids were then imported into RELION 4.0.1 (*34*). Capsids were re-extracted with 4× binning (4.272 Å/pixel) and reprocessed using CTFFIND4 (*35*) for CTF estimation. An initial 3D model was generated using I3 symmetry, followed by 3D refinement. To focus on the pentavalent capsomer, symmetry expansion was applied in I3, and capsomers were extracted at a fixed coordinate, resulting in 1,598,036 pentavalent capsomers. These were imported back into cryoSPARC using pyem and subjected to homogeneous refinement. Multiple rounds of heterogeneous refinement were performed to remove junk particles, yielding a final dataset of 239,460 pentavalent capsomers. C5 symmetry expansion was applied to each pentavalent capsomer, generating four additional symmetry-related orientations at rotations of 72°, 144°, 216°, and 288°. Subsequent 3D classification was performed with no imposed symmetry by using a focused mask on the central L2 density. This symmetry expansion increases the likelihood that equivalent L2 features are aligned during 3D classification, while releasing the fivefold symmetry averaging imposed by the icosahedral reconstruction. Multiple conformations of the L1/L2 complex at the pentavalent vertex were thereby resolved, and representative classes were subjected to further local refinement.

For analysis of hexavalent capsomers, the original particles were subjected to C5 symmetry expansion. The Volume Alignment tool in cryoSPARC was used to shift the extraction box to gain surrounding 7,990,103 hexavalent capsomers. Aligning the 3D map in cryoSPARC was then used to reorient particles along the central axis of the hexavalent capsomer. Similar heterogeneous refinement procedures are applied to generate 1,168,371 high-quality capsomers, which were further subjected to 3D classification and local refinement, enabling the reconstruction of distinct conformations L1/L2 complex at the hexavalent capsomers.

Movie stacks of L1-only PsV were processed using a similar strategy. After Topaz autopicking and 2D classification, 127,674 capsids were selected. From RELION processing, 1,182,505 pentavalent capsomers were extracted and locally refined with C5 symmetry. In parallel, 5,754,056 hexavalent capsomer particles were locally refined with C1 symmetry.

The same data collection and processing strategy was applied to the untagged L1/L2 PsV sample. 2,976 movie stacks were acquired, from which 41,908 capsids were auto-picked. Subsequent subparticle extraction yielded 502,863 pentavalent capsomers and 2,513,739 hexavalent capsomers. Comparable L2-associated densities were observed in both types of capsomers.

### Mass spectrometry

Mass spectrometry and preliminary data analysis was performed by Poochon Scientific (Frederick, MD, USA). Gradient-purified HPV16 L1/L2 and L1-only PsVs were concentrated in PBS and subjected to in-solution trypsin/lys-C digestion. The digested peptide mixture was then concentrated, desalted and reconstituted in 10 μl of 0.1% formic acid. Each 8 μl peptide sample was analyzed by a 110 min LC/MS/MS run using a Thermo Scientific Orbitrap Exploris 480 Mass Spectrometer and a Thermo Dionex UltiMate 3000 RSLCnano System. Digested peptides were loaded onto a peptide trap cartridge at a flow rate of 5 μL/min. Trapped peptides were eluted onto a reversed-phase Easy-Spray Column PepMap RSLC, C18, 2 μM, 100A, 75 μm × 250 mm (Thermo Scientific) using a linear gradient of acetonitrile (3-36%) in 0.1% formic acid. The elution duration was 60 min at a flow rate of 0.3 μl/min. Eluted peptides from the Easy-Spray column were ionized and sprayed into the mass spectrometer, using a Nano Easy-Spray Ion Source (Thermo) under the following settings: spray voltage, 1.6 kV, Capillary temperature, 275°C. Other settings were empirically determined.

Proteome Discoverer 3.1 software (Thermo, San Jose, CA) based on the SEQUEST algorithm were used to search raw data files against a sequence database containing HPV16 L1, HPV16 L2 with a C-terminal 3x FLAG tag, and human proteins. Peptide identification and label-free quantification were performed with peptide abundance values determined from MS1 precursor ion signals. For each identified protein, the average peptide intensity was calculated as the mean abundance of the three most abundant peptides assigned to that protein. Proteins were ranked according to their average peptide intensities. The relative abundances of the ten most abundant proteins in each PsV preparation were calculated as percentages of the summed average peptide intensities of these ten proteins. The resulting values were expressed as percentages and visualized as pie charts using Microsoft Excel (Microsoft Corporation, Redmond, WA).

The predicted copy numbers of HPV16 L1, L2, and histones per capsid were obtained from previous studies (*5, 36*). The copy numbers of L1, L2, histones, ferritin, and other contaminating proteins per capsid were estimated based on the relative average peptide intensities of the ten most abundant proteins obtained by label-free quantitative mass spectrometry. The average peptide intensity of each protein was normalized to that of HPV16 L1, assuming 360 copies of L1 per capsid. The normalized values were then multiplied by 360 to estimate the copy number of each protein per capsid.

### AlphaFold3 prediction of papillomavirus capsid protein structures

AlphaFold3 was used to predict structures of the L2 protein alone and within the L1 pentamer by using the AlphaFold Server (https://alphafoldserver.com) (*25*). Amino acid sequences of a single molecule of HPV16 L2, with inserted tags if indicated, and five molecules of HPV16 L1 were used as input for AlphaFold3 with default settings. Amino acid sequences of wild-type L1 and L2 proteins for HPV16, HPV5, BPV1, and CmPV1 were obtained from the PapillomaVirus Episteme (https://pave.niaid.nih.gov/). For the multiple independent predictions of L2 associated with the pentamer, AlphaFold3 was run repeatedly with identical amino acid sequences, with or without the auto-seed option, which generates distinct predictions for each run. Results were downloaded locally and predicted structures were visualized using PyMOL (The PyMOL Molecular Graphics System, Version 3.1.4.1, Schrödinger, LLC) or UCSF ChimeraX, Version 1.10.1 (*37*).

The full-length, wild-type HPV16 L1 protein was used in all AlphaFold3 modeling, unless noted otherwise. The following HPV16 L2 proteins were analyzed: wild-type, wild-type with a C-terminal 3xFLAG tag, or a mutant lacking the C-terminal 10 amino acids. For HPV5, BPV1, and CmPV1, wild-type L1 and L2 were used. Predictions were also generated for wild-type HPV16 L2 associated with HPV5 L1.

For defining the location and length of the L2 finger-like structure, the most distal part of the protruding finger relative to the L2 body was defined as the position of the finger. The length of the finger was defined as the number of amino acids comprising the finger segment.

To compare the variability of the structures of the various L2 elements, each segment of HPV16 L2 was defined based on its predicted structural characteristics: N-terminal helical region (residues 1-67), N-terminal disordered region (NDR, residues 68-223), central structured region (CSR, residues 224-332), C-terminal disordered region through the C-terminus (CDR+, residues 333-473). Three AlphaFold3 prediction models were compared pairwise to each other, and intact L2 or individual segments were aligned and RMSD values calculated using the matchmaker tool in ChimeraX. To create density models of AlphaFold3 predictions, 100 models were combined into a density map by using ChimeraX molmap. AlphaFold3 models were aligned using the matchmaker tool with default settings and a density model was created for L2 using the molmap command across the entirety of L2. The threshold level was adjusted using the volume viewer tool in ChimeraX. Individual threshold levels are noted in each figure as the sigma value for the chosen threshold level, calculated as the contour level divided by the root-mean-square (RMS).

The electrostatic surface potentials of L1 and L2 were calculated and visualized using UCSF ChimeraX Version 1.10.1, based on the cryo-EM structure model of L1 determined in this study and an AlphaFold3 predicted structure of L2 with a C-terminus finger. Molecular surfaces were colored according to the calculated electrostatic potential, with negatively charged regions shown in red, neutral regions in white, and positively charged regions in blue.

Interactions between L1 and L2 were counted in the AlphaFold3-generated PDB structures with MDtraj using a 4.5 Å cutoff between heavy atoms (*38*). Results were aggregated for 40 models with NDR fingers and 46 models with CDR fingers.

### Molecular dynamics

All-atom molecular dynamics (MD) simulations were performed on the Yale Bouchet High Performance Computer with GROMACS v2024.6 parameters and topologies generated by the CHARMM-GUI Solution Builder (*39–42*). The system used the CHARMM36m force-field due to its ability to accurately model disordered protein dynamics (*43*). The starting conformations of L1 and L2 were obtained from the AlphaFold3 modeling performed in this study. CHARMM-GUI was used to truncate L1 to residues 1-480, thereby removing the disordered C-terminal tails, and to introduce non-standard protonation at specific residues. For the simulations, L1 residues Glu167, His259, Glu362, and His366 were protonated because of their elevated local pKa’s as predicted by H++ webserver (http://biophysics.cs.vt.edu/H++) (*44*). All other titratable residues in L1 and all residues in L2 were assigned their standard protonation states at pH 7.4.

All systems were solvated in a 150 mM neutralizing NaCl cubic water box with dimensions based on the size of each model (edges extending 30 Å from the protein[s]) by using the TIP3 water model (*45*). Each replicate simulation followed the standard CHARMM-GUI energy minimization, NVT equilibration, and NPT production protocols. Production runs were performed without position restraints at 1 bar and a temperature of 310.15 K maintained by a C-rescale barostat and V-rescale thermostat. All simulations were run for at least 500 ns with a 2 or 2.5 fs timestep with two replicates per condition, for a total simulation time of ≥1 µs for each simulation.

Analysis of trajectories was performed using the GROMACS analysis toolkit and MDTraj (*38*), with visualization and modeling performed using VMD version 1.9.4 (*46*). RMSD and RMSF calculations were performed using backbone Cα’s. RMSF estimated density distributions were determined by pooling RMSF measurements for each residue across replicate simulations and using a Gaussian Kernel Density Estimation function in SciPy. L2 residues inside and beneath the L1 chamber/pore were estimated visually for the simulation shown in Fig. 2G, with the following L2 residues being considered inside: 98-143, 162-194, and 395-473, and the following residues considered beneath: 1-97, 144-161, 195-394.

### Co-Immunoprecipitation assay

Four µL (∼0.35 μg of L1) of PsV sample containing L2 with a C-terminal HiBiT tag was resuspended in 300 µL of 50 mM HEPES (pH 7.4), 150 mM NaCl, 0.1% Triton X-100, and 1X protease inhibitor cocktail (from tablets, Roche,11873580001) in PBS. A 30 µL aliquot was reserved as the input sample, while the remaining 270 µL was incubated overnight at 4°C in the absence of antibody or with anti-HiBiT mouse primary antibody (1:200 dilution; Promega, catalogue # N7200). Forty µL of pre-calibrated Protein G-conjugated Dynabeads (Thermofisher, catalogue # 10003D) was then added to the mixture and incubated for an additional hour at 4°C. The beads were then sequestered using a magnetic rack, and 30 µL of the resulting supernatant was collected as the unbound fraction. The beads were washed four times with PBST [PBS containing 0.1% Tween-20 (Sigma, Catalogue # P7949). Bound proteins were eluted by heating the beads at 100°C for 7 minutes in 60 µL of 2X Laemmli sample buffer supplemented with β-mercaptoethanol. The input, eluate, and supernatant fractions were analyzed by SDS-polyacrylamide gel electrophoresis and western blotting with antibodies recognizing L1 (HRP CAMVIR, Santa Cruz Biotechnology, Catalogue # SC-47699, 1:1000 dilution) or L2 (anti-HiBiT antibody; Promega, catalogue # N7200, 1:1000 dilution).

### Split luciferase assay

A split luciferase assay was performed to quantitatively measure exposure of the L2 C-terminus. Density gradient-purified HPV16 PsVs lacking a HiBiT tag or containing a HiBiT tag appended to the C-terminus of L2 were used. To generate a source of LgBiT, we first inserted the LgBiT coding sequence in-frame upstream of the eGFP coding sequence in pEGFP-2xFYVE (Addgen plasmid #140047) to construct pLgBiT-eGFP-2xFYVE, which was used to transfect 293TT cells. 48 h later, cells were harvested by scraping from the culture dish in PBS and stored at −80°C. Prior to use, the cells were thawed and sonicated five times for 2 s at 75% power to generate the crude cell lysate containing LgBiT. To evaluate L2 C-terminal exposure on intact PsVs, 0.25 μL of Nano-Glo Luciferase Assay substrate (Promega, Catalogue # N1110) and 100 μL of lysate containing LgBiT were added to ∼0.5 μg PsV per well in a 96-well assay plate (Corning Costar, 3903). Luminescence signals were measured at room temperature by using a GloMax plate reader every min for 100 reads. To show that activity was due to L2 associated with L1, PsVs were incubated with 2 μL anti-L1 antibody (V5, gift of Neil Chistensen) or non-specific IgG1 control antibody (BD Biosciences, Catalogue # 555746) at 4°C for 30 min, followed by incubation with 20 μL of pre-calibrated Protein G-conjugated Dynabeads (Thermo Fisher Scientific, catalogue # 10003D) at 4°C for 1 h. Bead-bound and supernatant fractions were separated using a magnetic rack, and the supernatant fraction was collected for analysis. To determine the total amount of luciferase activity due to HiBiT-tagged L2 incorporated into the capsid, PsVs were heated at 99°C for 10 min to dissociate the capsids prior to assay.

### Immunogold staining and transmission electron microscopy

PsV samples were filtered out from OptiPrep into PBS and concentrated approximately 2 to 4-fold by using a 100 kDa cutoff cellulose filter (Millipore, Sigma). All subsequent steps were carried out at room temperature. Formvar grids (Carbon, 400-mesh; Ted Pella, Inc., Redding, California) were glow-discharged using a PELCO easiglow instrument and then floated on 15 µL droplets of concentrated PsV sample (∼3 µg PsV) for 8–10 minutes. Following two washes with 30 µL droplets of filtered PBS (pH 7.0), the grids were blocked on 30 µL droplets of 0.5%–1% bovine serum albumin for 30 minutes. The grids were then incubated on 15 µL droplets containing primary antibody recognizing an epitope tag for one hour at room temperature, washed four times with PBS, and then incubated on 15 µL droplets of 10 nm gold-488AF conjugated goat-anti-mouse secondary antibody (1:10 dilution, Thermofisher, catalogue # A-31561) for 30 minutes. Anti-HA (Cell Signaling, catalogue # C29F4) and anti-HiBiT (1:5 dilution, Promega, # N7200) were used as primary antibodies. After four additional washes with 30 µL droplets of filtered PBS (pH 7.0), the samples were negatively stained with 2% pH-neutralized uranyl formate and imaged using a JEOL JEM-1400 transmission electron microscope.

### Neutralization experiments

For neutralization experiments, 1×10^5^ HeLa S3 cells per well were seeded in 12 well plates and incubated overnight. PsV contained the HcRed reporter plasmid and untagged L2 or L2 with a HiBiT tag at position 385 or a HiBiT or HA tag at C-terminus. PsV (∼0.1 μg) was incubated in DMEM in the absence of antibody or with 2 μL of anti-HiBiT antibody (Promega, catalogue # N7200) or anti-HA antibody (Cell Signaling, catalogue # C29F4) for 1 h at room temperature prior to addition to cells at a multiplicity of infection (MOI) of ∼1. 48 h post infection (hpi), the cells were analyzed by flow cytometry by using CytoFLEX LX flow cytometer (Beckman Coulter, Brea, CA).

### Western blotting

Various amounts of untagged HPV16 PsV or PsV containing a HiBiT tag in L2 at position 385 or the C-terminus were subjected to SDS-polyacrylamide gel electrophoresis and western blotting with an antibody recognizing the HiBiT tag (anti-HiBiT, 1:5000 dilution; Promega, catalogue # N7200) or L2 itself (anti-HPV16 L2 [2JGmab#5], 1:5000 dilution; Santa Cruz, catalogue # sc-65709). Goat-anti-mouse-HRP (Jackson Immuno, cat#115--035-146) was used as secondary antibody at 1:10,000 dilution.

## Acknowledgments

We thank Yuka Takeo for the p16SheLL plasmid expressing only L1, and we thank Neil Christensen for the V5 anti-L1 antibody. This work was supported by grants from the National Institutes of Health to DD (R35CA242462) and to JL (GM124378 and GM110243). Cryo-EM data were collected at the Yale CryoEM Resource. Initial sample screening was performed with Glacios microscopes at Yale Science Hill and Yale School of Medicine, and single-particle cryo-EM data were collected with the Krios microscope at Yale West Campus. Flow cytometry was conducted at the Yale Flow Cytometry Shared Resource supported in part by the Yale Cancer Center Support Grant (P30CA016359).

## Author Contributions

**Conceptualization:** CO, HY, JL, DD

**Investigation:** CO, HY, PB, FH, KM, JC

**Writing original draft:** CO, HY

**Revisions:** all authors

**Supervision:** DD and JL

**Funding acquisition:** DD and JL

## Data, code, and materials availability

HPV16 PsV cryo-EM maps have been deposited in the Electron Microscopy Data Bank (EMDB) and Protein Data Bank (PDB) with the following accession codes: HPV16 L1-only PsV pentavalent capsomers (PDB ID 37YL, EMD-78644); HPV16 L1-only hexavalent capsomers (PDB ID 37YM, EMD-78645); HPV16 L1/L2FLAG PsV pentavalent capsomers (PDB ID 37YB, EMD-78635); HPV16 L1/L2FLAG PsV hexavalent capsomers (PDB ID 37YJ, EMD-78643); HPV16 L1/L2 (untagged) PsV pentavalent capsomers (PDB ID 37YN, EMD-78646); HPV16 L1/L2 (untagged) PsV hexavalent capsomers (PDB ID 37YO, EMD-78647). No new code was generated for this study. All materials are available from the authors upon request.

**Supplemental Table 1.** Metrics of cryo-EM analysis of HPV16 pseudovirus particles.

|  | Pentavalent<br>capsomer of<br>PsV<br>L1/L2FLAG | Hexavalent<br>capsomer of<br>PsV<br>L1/L2FLAG | Pentavalent<br>capsomer of<br>PsV L1-only | Hexavalent<br>capsomer of<br>PsV L1-only | Pentavalent<br>capsomer of<br>PsV L1/L2<br>(untagged) | Hexavalent<br>capsomer of<br>PsV L1/L2<br>(untagged) |
| --- | --- | --- | --- | --- | --- | --- |
| **Data collection** |  |  |  |  |  |  |
| Magnification | ×81,000 |  |  |  |  |  |
| Voltage (kV) | 300 |  |  |  |  |  |
| Total electron dose (e<sup>-</sup>/Å<sup>2</sup>) | 55 |  |  |  |  |  |
| Defocus range (μm) | -1.6 to -2.2 |  |  |  |  |  |
| Pixel size (Å) | 1.068 |  |  |  |  |  |
| Micrographs (no.) | 3,660 |  | 3,594 |  | 2,976 |  |
| **Data processing** |  |  |  |  |  |  |
| Symmetry imposed | C5 | C5 | C5 | C5 | C5 | C5 |
| Final particle images (no. ) | 1,598,036 | 7,990,103 | 1,150,919 | 5,754,056 | 502,863 | 2,513,739 |
| Map resolution (Å) 0.143 FSC | 2.16 | 2.37 | 2.28 | 2.24 | 2.37 | 2.36 |
| Map sharpening B factor (Å<sup>2</sup>) | -64.8 | -72.6 | -82.6 | -92.5 | -74.6 | -64.8 |
| **Refinement** |  |  |  |  |  |  |
| CC (model vs. data) | 0.82 | 0.87 | 0.86 | 0.90 | 0.86 | 0.87 |
| Chain count | 5 | 5 | 5 | 5 | 5 | 5 |
| Non-hydrogen atoms | 17,000 | 17,000 | 17,000 | 17,000 | 17,000 | 17,000 |
| Protein residues | 2,160 | 2,160 | 2,160 | 2,160 | 2,160 | 2,160 |
| Ligands | 0 | 0 | 0 | 0 | 0 | 0 |
| B factors (Å<sup>2</sup>) |  |  |  |  |  |  |
| Proteins | 40.22 | 54.65 | 35.67 | 51.92 | 39.17 | 37.50 |
| Ligands | / | / | / | / | / | / |
| R.m.s. deviations |  |  |  |  |  |  |
| Bond lengths (Å) | 0.004 (0) | 0.003 (0) | 0.003 (0) | 0.003 (0) | 0.003 (0) | 0.004 (0) |
| Bond angles (°) | 0.672 (2) | 0.632 (2) | 0.587 (5) | 0.575(0) | 0.590 (1) | 0.575 (0) |
| **Validation** |  |  |  |  |  |  |
| MolProbity score | 1.93 | 1.73 | 1.70 | 1.76 | 1.86 | 1.82 |
| Clashscore | 8.00 | 6.75 | 6.46 | 6.22 | 6.46 | 6.04 |
| **Ramachandran plot** |  |  |  |  |  |  |
| Favoured (%) | 96.68 | 96.50 | 96.92 | 96.92 | 96.64 | 96.36 |
| Allowed (%) | 3.32 | 3.50 | 3.08 | 3.08 | 3.36 | 3.64 |
| Outliers (%) | 0.00 | 0.00 | 0.00 | 0.00 | 0.00 | 0.00 |
| PDB code | 37YB | 37YJ | 37YL | 37YM | 37YN | 37YO |
| EMDB code | EMD-78635 | EMD-78643 | EMD-78644 | EMD-78645 | EMD-78646 | EMD-78647 |

## Supplemental Figure legends

**Fig. S1.**
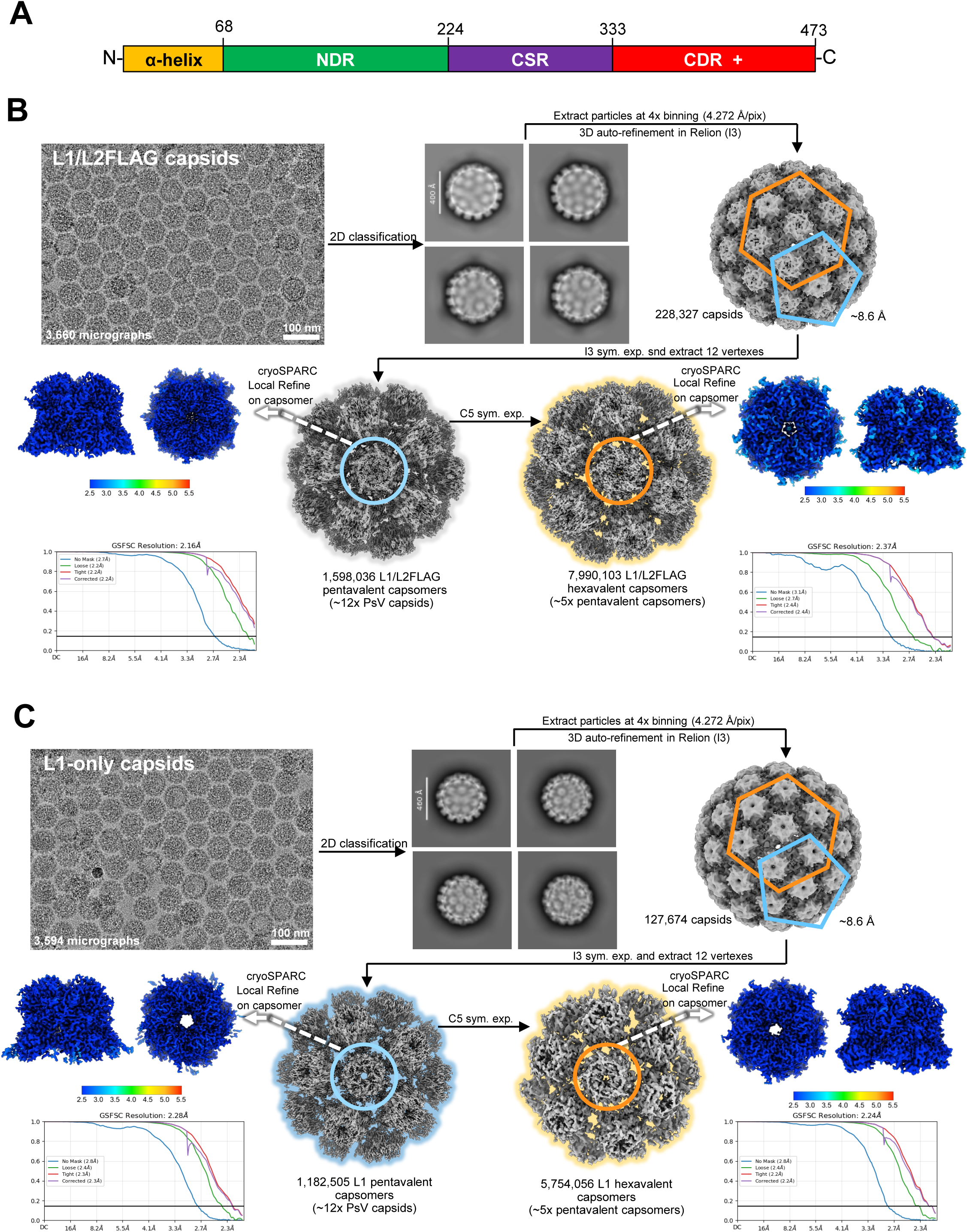
Cryo-EM data processing workflow for HPV16 L1-only and L1/L2 capsids and organization of L2 proteins. A. Cartoon showing overall organization of the L2 protein. Numbers indicate amino acid positions in HPV16 L2. The C-terminal disordered region and downstream sequences to the C-terminus of L2 are designated C-term disordered +. B,C. Representative micrographs, 2D class averages, and 3D reconstruction workflows for L1/L2FLAG (B) and L1-only (C) PsV, respectively. After capsid 3D refinement, symmetry expansion enabled extraction of pentavalent (blue pentagons and circles) and hexavalent (orange hexagons and circles) capsomers from both datasets. In the L1/L2 dataset, local refinement produced high-resolution maps for capsomers (2.16 Å for pentavalent and 2.37 Å for hexavalent). Local refinement produced high-resolution maps for L1-only capsomers (2.38 Å for pentavalent and 2.59 Å for hexavalent). Subsequent 3D classification and refinement revealed multiple conformations at both vertex types, reflecting structural heterogeneity introduced by L2. Final reconstructions and FSC curves are shown for each capsomer type. Local resolution is also estimated in cryoSPARC and is shown using a gradient color representation.

**Figure S2.**
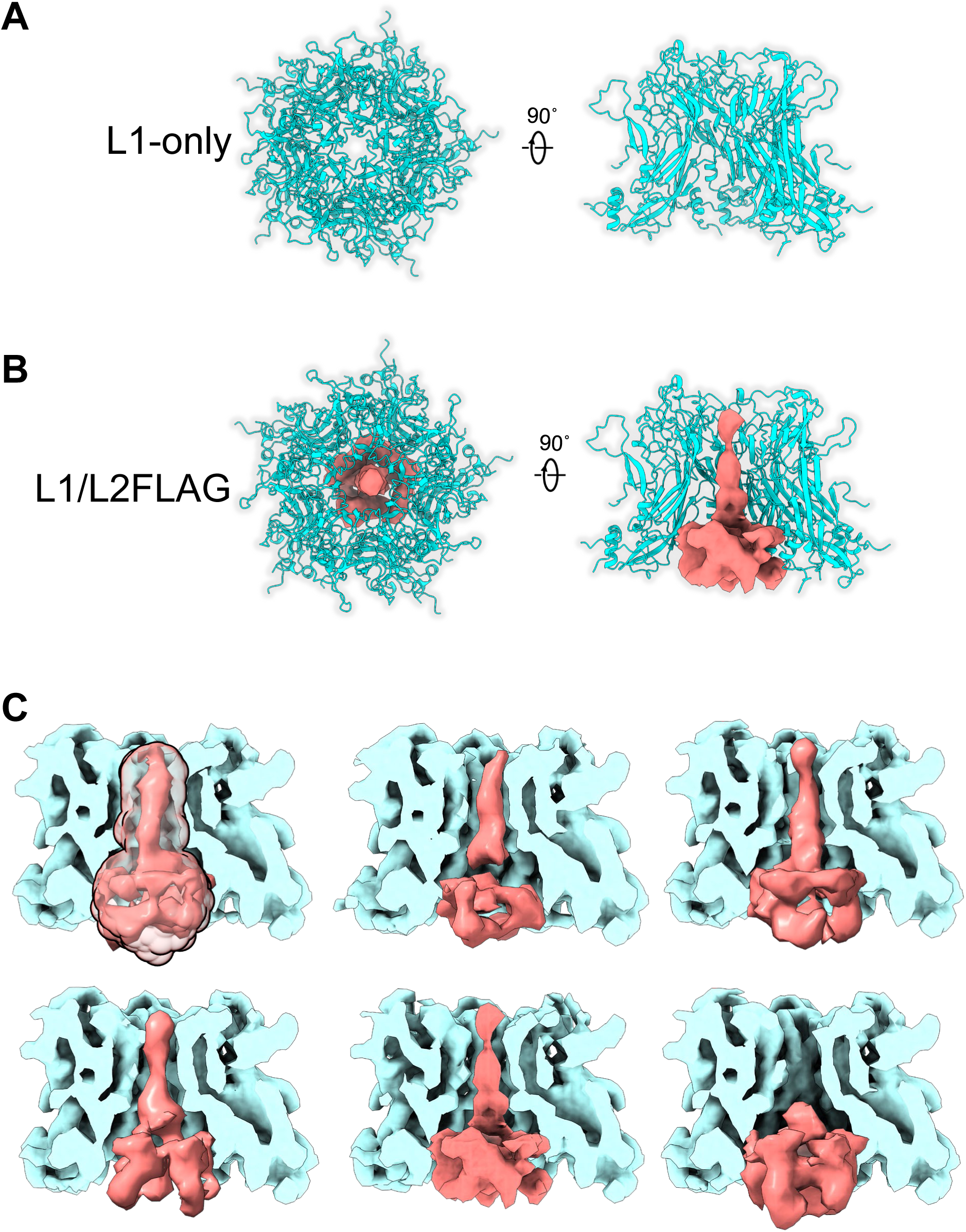
Cryo-EM analysis of hexavalent capsomers. A. Top view (left image) and side view (with the front two L1 monomers hidden from view for clarity, right image) of hexavalent capsomers from L1-only capsids, shown in ribbon representation. B. Top and side views of L1/L2FLAG hexavalent capsomers as in panel A. Potential L2 density in pink is represented by one of the selected 3D classes. C. Representative 3D classifications focused on the central densities within the pore and the L1 pentamer chamber in pentavalent capsomers in the L1/L2FLAG capsids. The transparent density in the upper left image indicates the mask region used for 3D classification. Side views are shown, with L1 in cross section.

**Figure S3.**
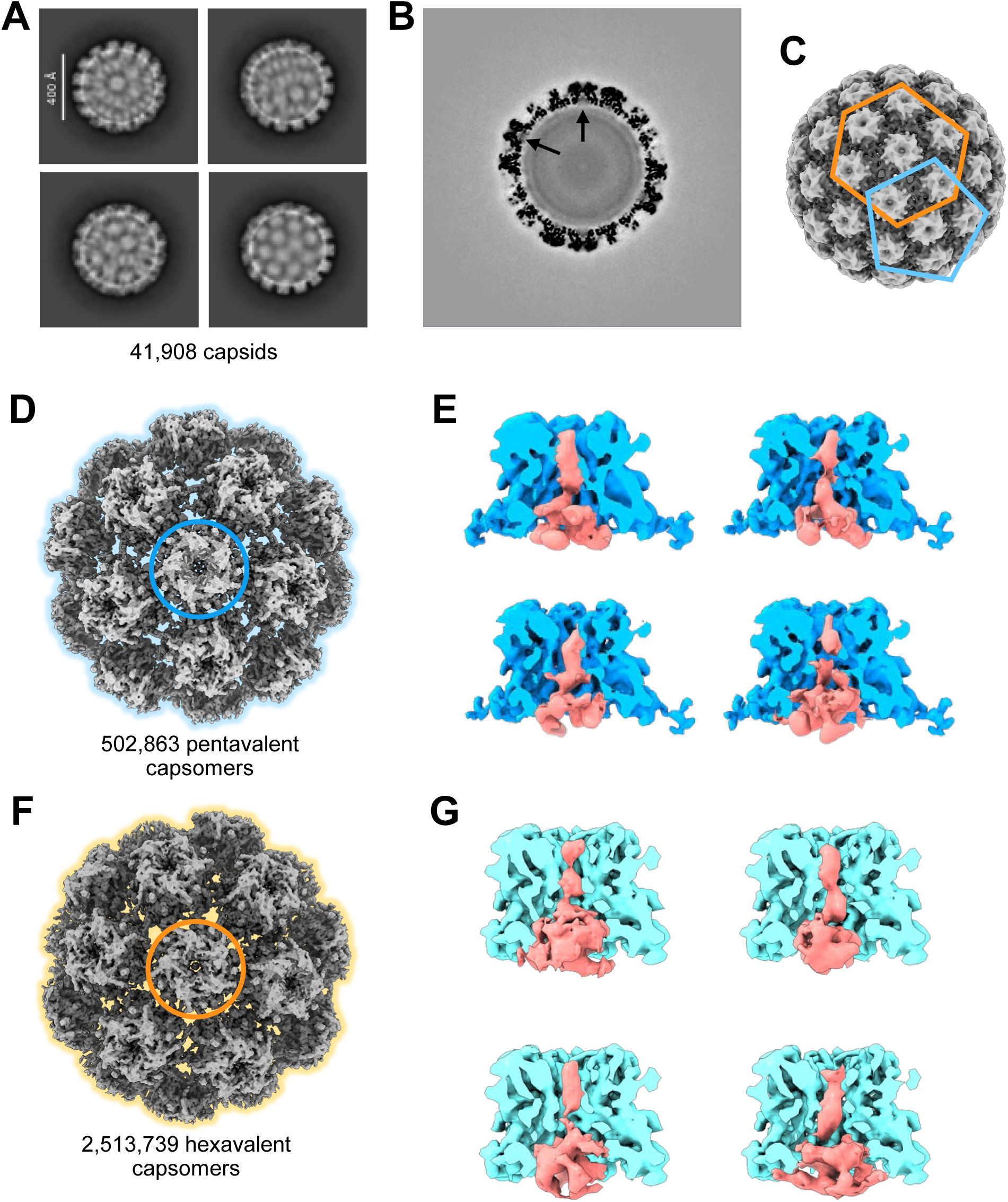
Cryo-EM analysis of capsids containing untagged L2. (A) Representative 2D class averages of capsid particles. (B) Central section through the capsid reconstruction generated with icosahedral symmetry. Arrows indicate densities corresponding to L2 in the pentamer chamber. (C) 3D view of the overall icosahedral capsid reconstruction. Pentavalent and hexavalent capsomers, outlined in blue and orange, respectively, were analyzed individually. (D) Focused 3D classification of pentavalent capsomer. (E) Cross-sectional view of four representative 3D classes showing L2 density (pink) within the pentavalent L1 capsomer (dark blue). (F) Focused 3D classification of hexavalent capsomer. (G) Cross-sectional view of four representative 3D classes showing L2 density (pink) within the hexavalent capsomer (light blue). In (D) and (F), the small central circles indicate the capsomers selected for focused analysis.

**Figure S4.**
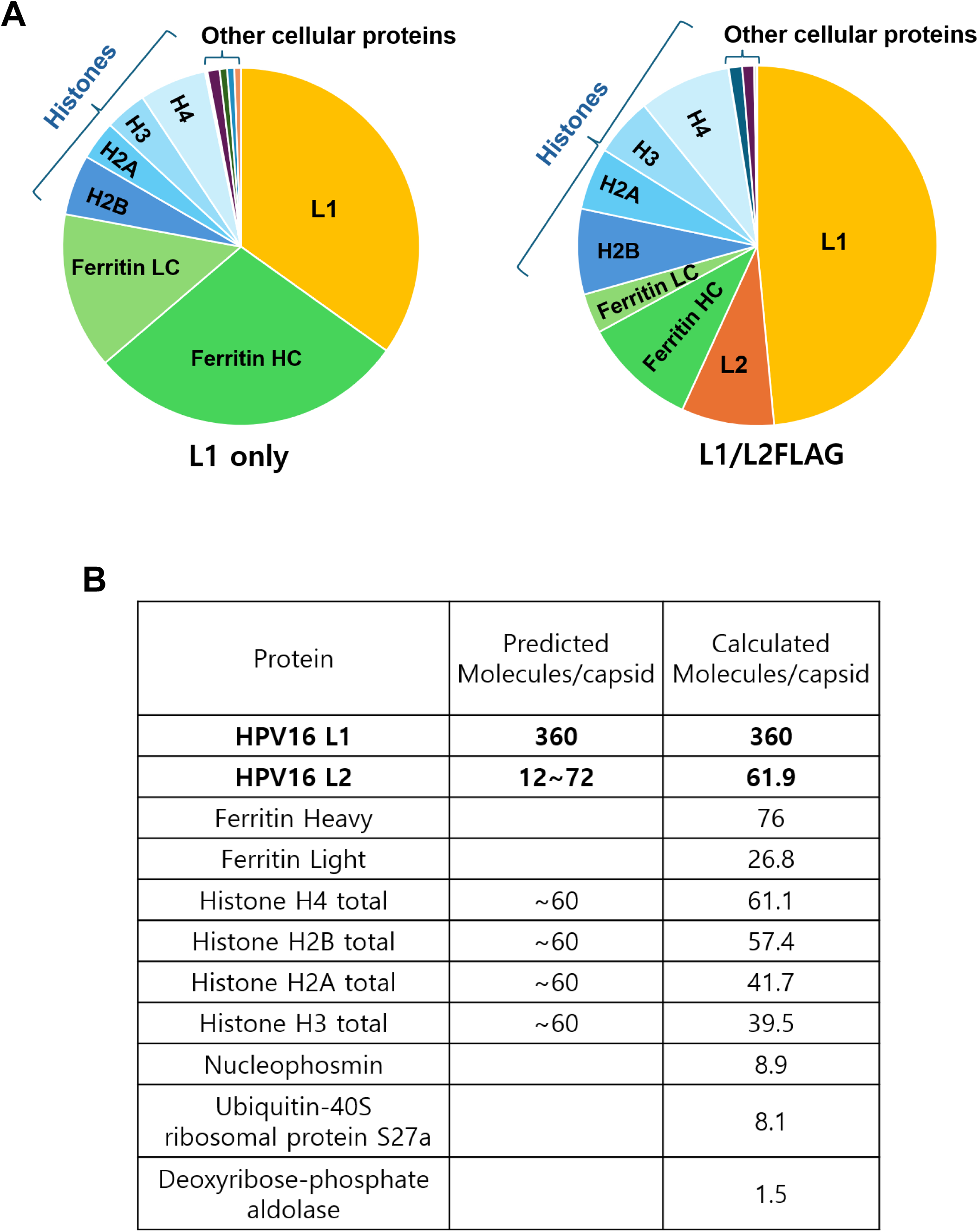
Mass Spectrometry analysis of pseudovirus preparations. A. Purified preparations of L1-only (left panel) and L1/L2FLAG (right panel) PsVs were subjected to mass spectrometry, and the abundance of proteins was estimated based on the relative abundance of the three most abundant peptides for each protein. B. Table showing the predicted number of molecules of the indicated proteins in each L1/L2FLAG PsV capsid, compared to the approximate number in the capsids calculated from on the mass spectrometry analysis. Nucleophosmin, Ubiquitin-40S ribosomal protein S27a, and Deoxyribose-phosphate aldolase were the most abundant contaminating proteins in the L1/L2FLAG PsV other than ferritin heavy chain (HC) and ferritin light chain (LC), which were more abundant in the L1-only PsV preparation. To predict the number of histone molecules in the capsid, each capsid is assumed to contain 30 nucleosomes (*36, 47*).

**Figure S5.**
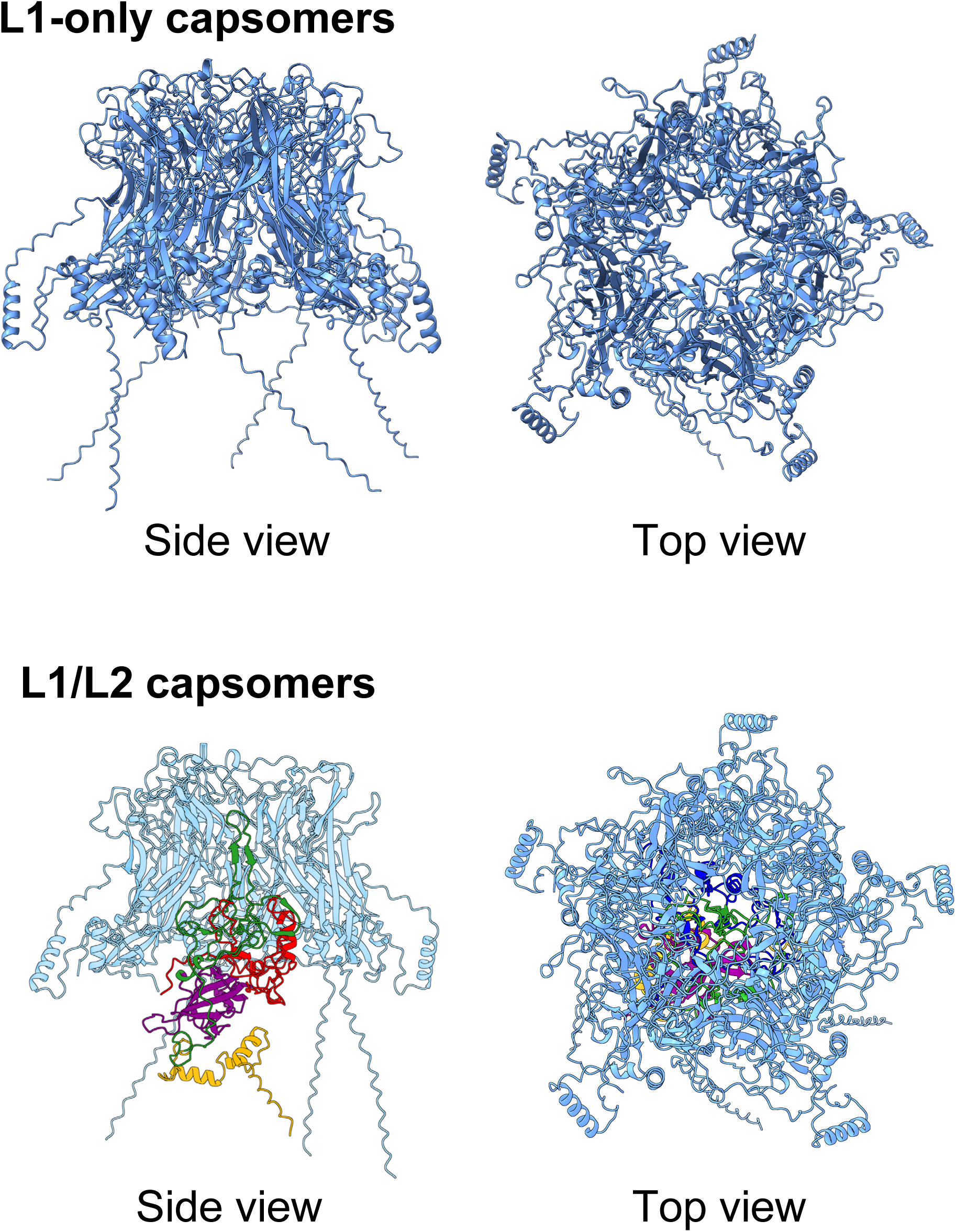
AlphaFold3 models of L1 pentamers in the presence and absence of L2. AlphaFold prediction of L1 only and L1/L2 were generated using five molecules of HPV16 L1, either without L2 (L1-only) or in the presence of a single molecule of HPV16 L2 (L1/L2). L1 pentamer are shown in blue, and L2 is color coded according to L2 domains, as shown in Figure S1A. For the side view of the L1/L2 capsomer, the two front L1 monomers were hidden from view to visualize L2.

**Figure S6.**
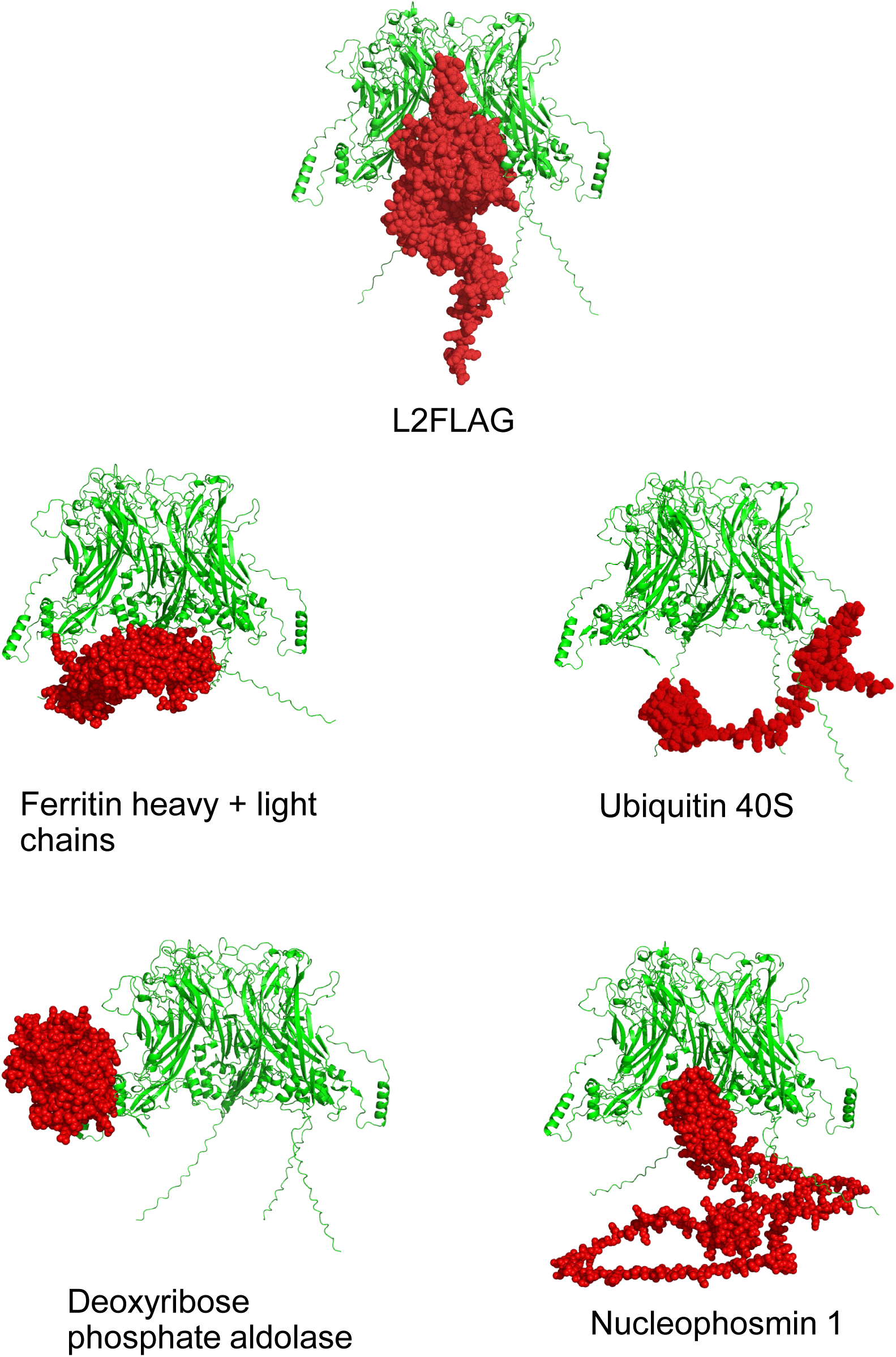
Contaminants in pseudovirus preparations are not predicted to align with the cryo-EM density. The most abundant non-histone cellular proteins identified in the L1/L2 PsV preparation by mass spectrometry were subjected to AlphaFold3 modeling with five molecules of HPV16 L1. A representative AlphaFold3 model of HPV16 L2 and the L1 pentamer is shown at the top. Capsomers are shown in side view with the front two L1 monomers hidden from view to visualize L2.

**Figure S7.**
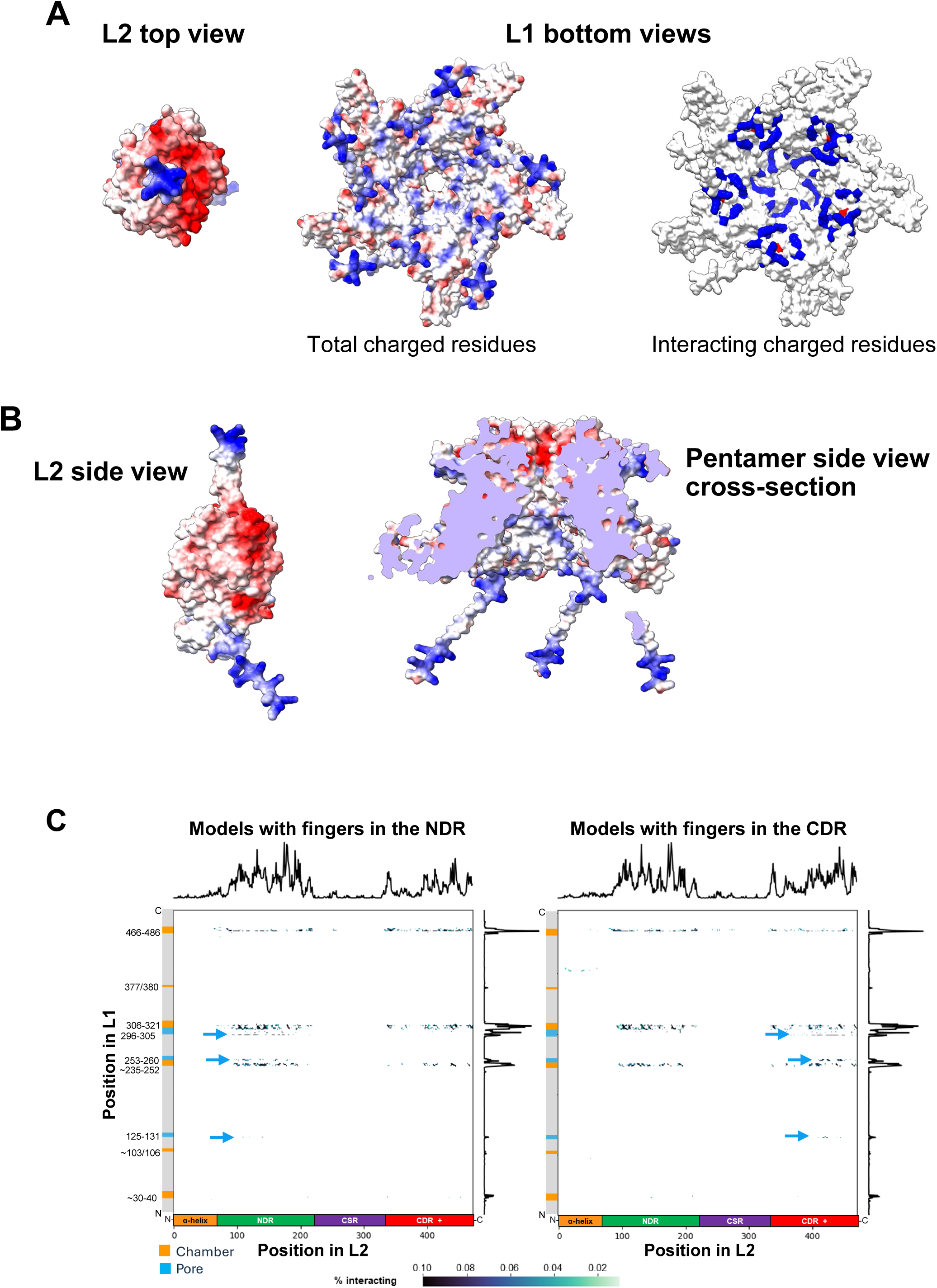
Analysis of the predicted L1/L2 interface and charge distribution in capsomers. A. Charge distribution of a representative AlphaFold3 model of the HPV16 L1 pentamer associated with L2 containing a C-terminal finger. Positively charged (basic) residues are colored blue and negatively charged (acidic) residues are colored red in these surface models. L2 model is presented in a view from the top (left image). The pentamer is presented in a view from the bottom, showing all charged surface residues (middle image) and charged residues that contact L2 (right image). B. Charge distribution as in panel A of a side view of L2 containing a C-terminal finger and a side view of a cross-section through the L1 pentamer. C. Two-dimensional heat map of interacting L1 and L2 residues. Cartoons of L1 (left side of each map) and L2 (bottom) of each map are schematically shown with various L1 and L2 segments color-coded. Residues lining the pentamer pore are shown in light blue, and residues lining the inner surface of the pentamer chamber are shown in orange. The aggregate results of 40 models with NDR fingers (left graph) and 46 models with CDR fingers (right graph) are shown. The frequency of interactions (i.e. heavy atoms of residues of L1 and L2 within 4.5 Å) is plotted in black lines on the top and right side of each graph. Individual interacting residue pairs are shown as dots, with more frequent interactions shown in darker blue (maximum intensity when 10% or more of the models show an interaction), as indicated by scale on bottom. The light blue arrows highlight interactions that occur within the pentamer pore.

**Figure S8.**
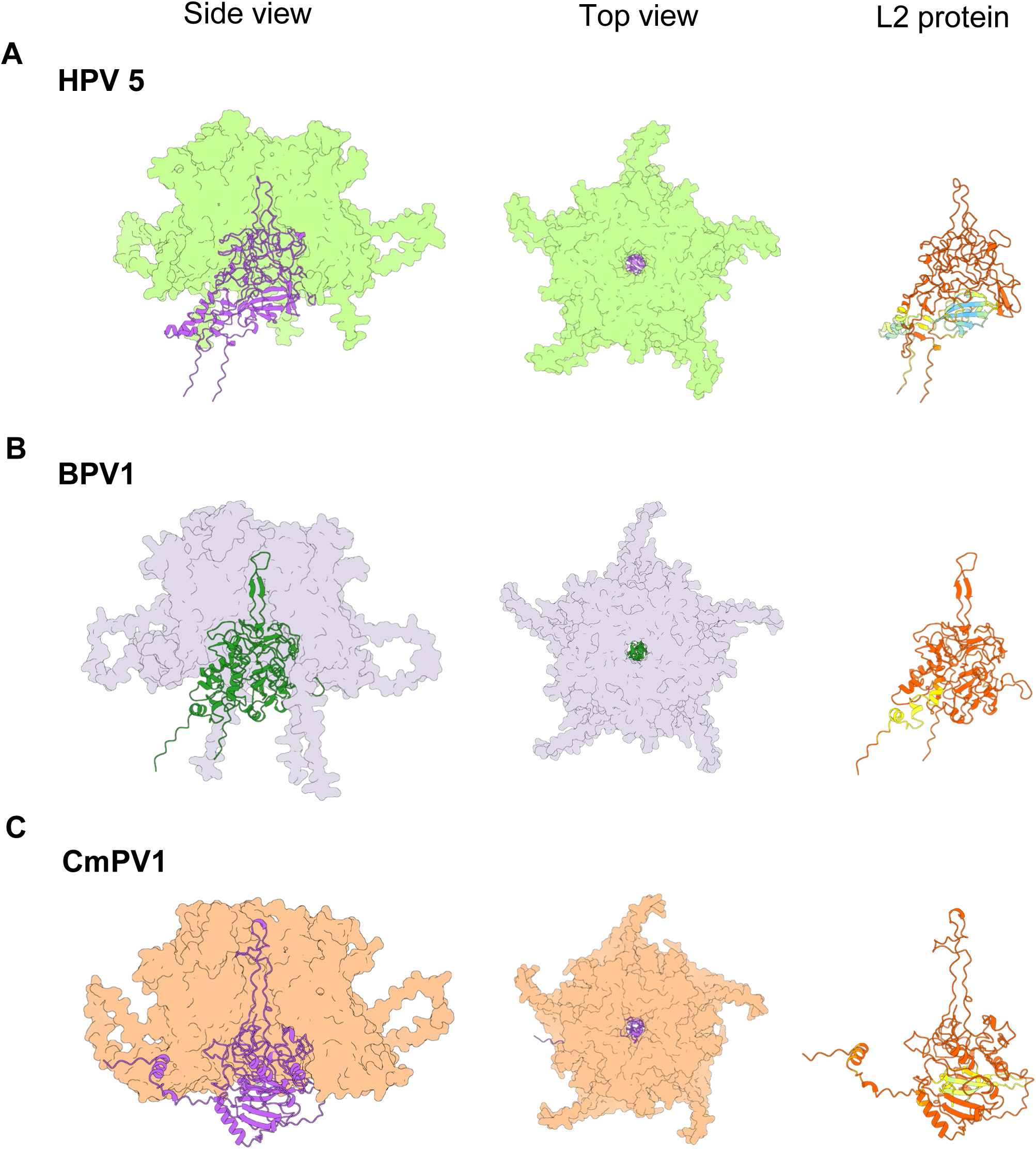
Divergent papillomaviruses are predicted to have similar L2 structures in the capsid. AlphaFold3 predictions of L1 pentamer in the presence of L2 from HPV 5, BPV1, and CmPV1 were generated using five molecules of the designated L1 protein and one molecule of the cognate L2 protein. L1 is shown in space-filling representation, and L2 is shown in ribbon form in the side (left) and top (middle) view of the capsomer. For the side view of the capsomer, the two front L1 molecules were hidden from view to visualize L2. In the isolated L2 protein view (right), L2 is color coded by pLDDT confidence scores, with the color scale shown in Figure 2C.

**Figure S9.**
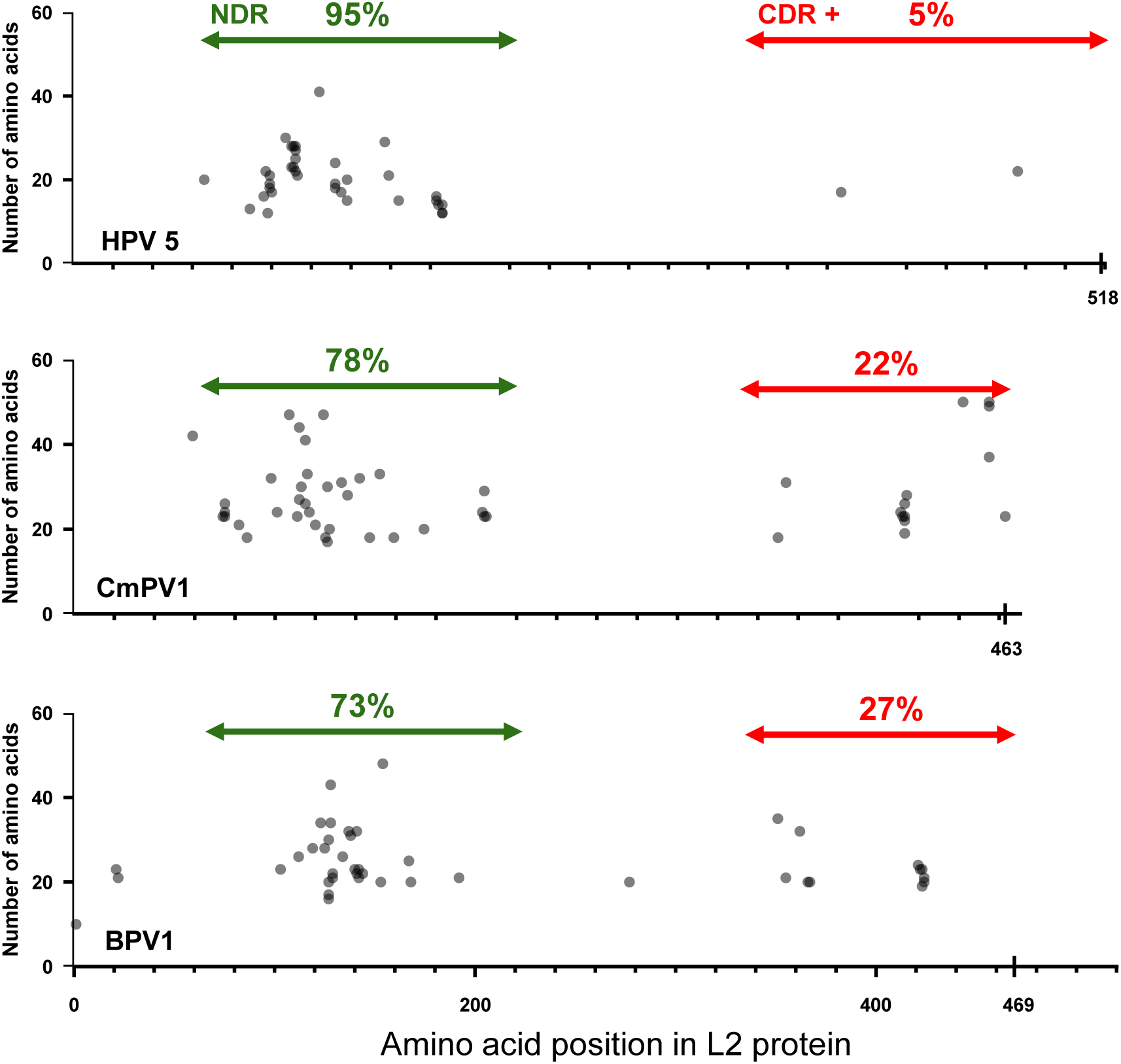
Divergent papillomaviruses display fingers distributed across disordered regions. Location of finger in AlphaFold3 models of HPV5, CmPV1, and BPV1. The position of the fingertip from multiple AlphaFold3 predictions of each virus are plotted as in Figure 3B.

**Figure S10.**
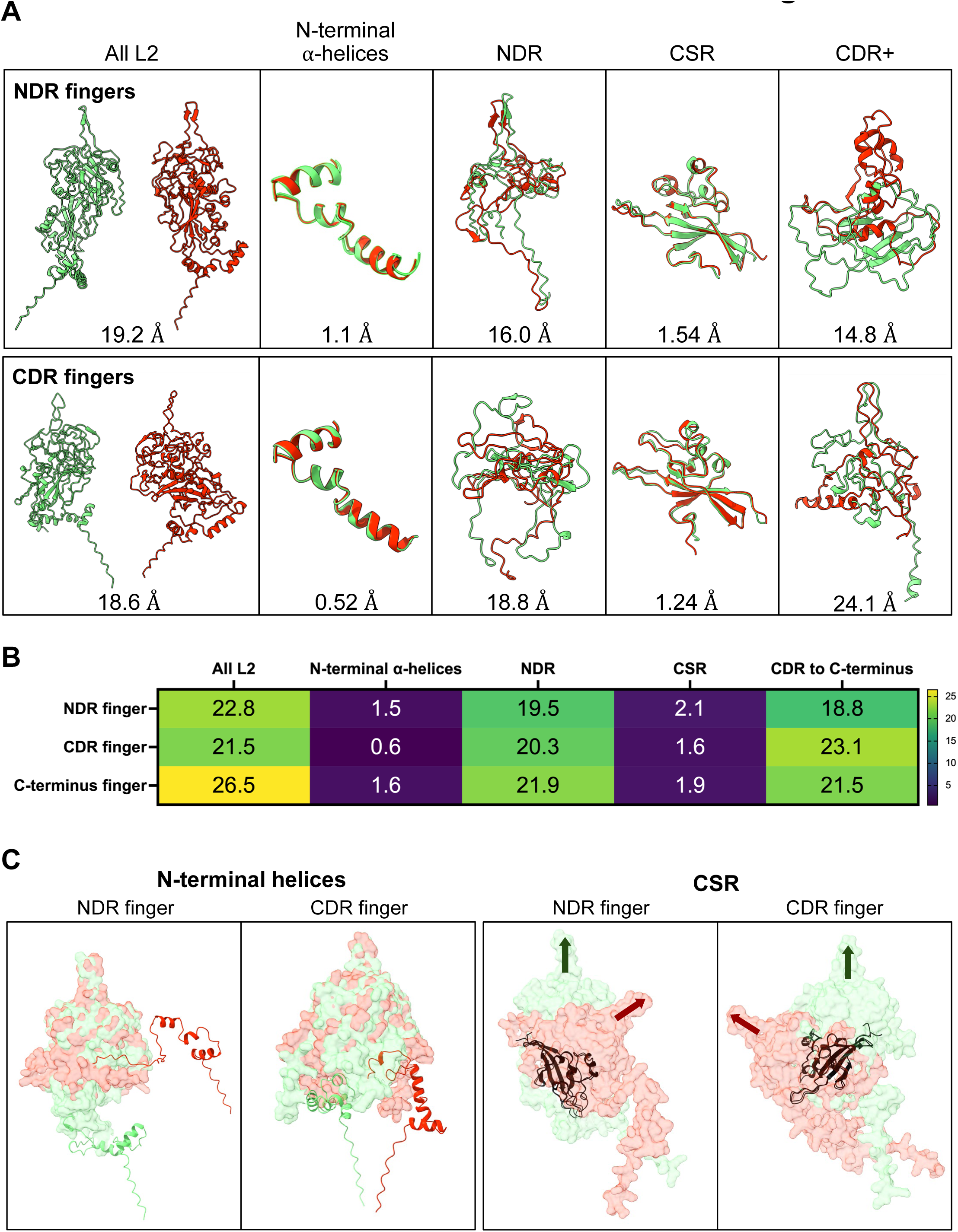
AlphaFold3 predicts high flexibility and mobility of different segments of L2 in capsomers. A. AlphaFold3 models with fingers from either the CDR or NDR (three for each) were compared across the entirety of L2 or for individual segments, namely amino acid positions 14 to 53, encompassing much of the N-terminal *a*-helices; positions 74 to 221, approximating the NDR; positions 222 to 334, corresponding to the CSR; and positions 335 to 473, the CDR+. A representative pair of models were selected for visualization, one shown in green and the other red. Top shows comparisons for two models with an NDR finger of similar length centered at position 179 and bottom shows comparisons for two models with a CDR finger of similar length centered at position 430. The segments compared are shown side by side in the case of full-length L2 or overlayed for the individual segments. The corresponding RMSD values are shown below each comparison. B. AlphaFold3 predictions for models with L2 fingers of similar lengths at position 179, 430, or the C-terminus were aligned across different segments of L2 and compared pairwise as in panel A (three models were compared for each set). Average RMSD values were calculated and presented for the indicated segment of L2, shown as a heat map with scale on right. C. The orientation of the N-terminal helices and the CSR were compared for representative AlphaFold3 predictions with L2 fingers centered in the NDR or CDR. Left two panels, the fingers and bodies of L2 were aligned, to highlight the different orientations of the N-terminal *a*-helices, shown as dark red and green ribbons with the remainder of L2 shown as light red and green molecular surfaces. For the right two panels, the β-sheets were aligned and depicted in black in ribbon representation. The remainder of L2 is shown as molecular surfaces with fingers indicated with arrows to highlight the different orientation of the β-sheet.

**Figure S11.**
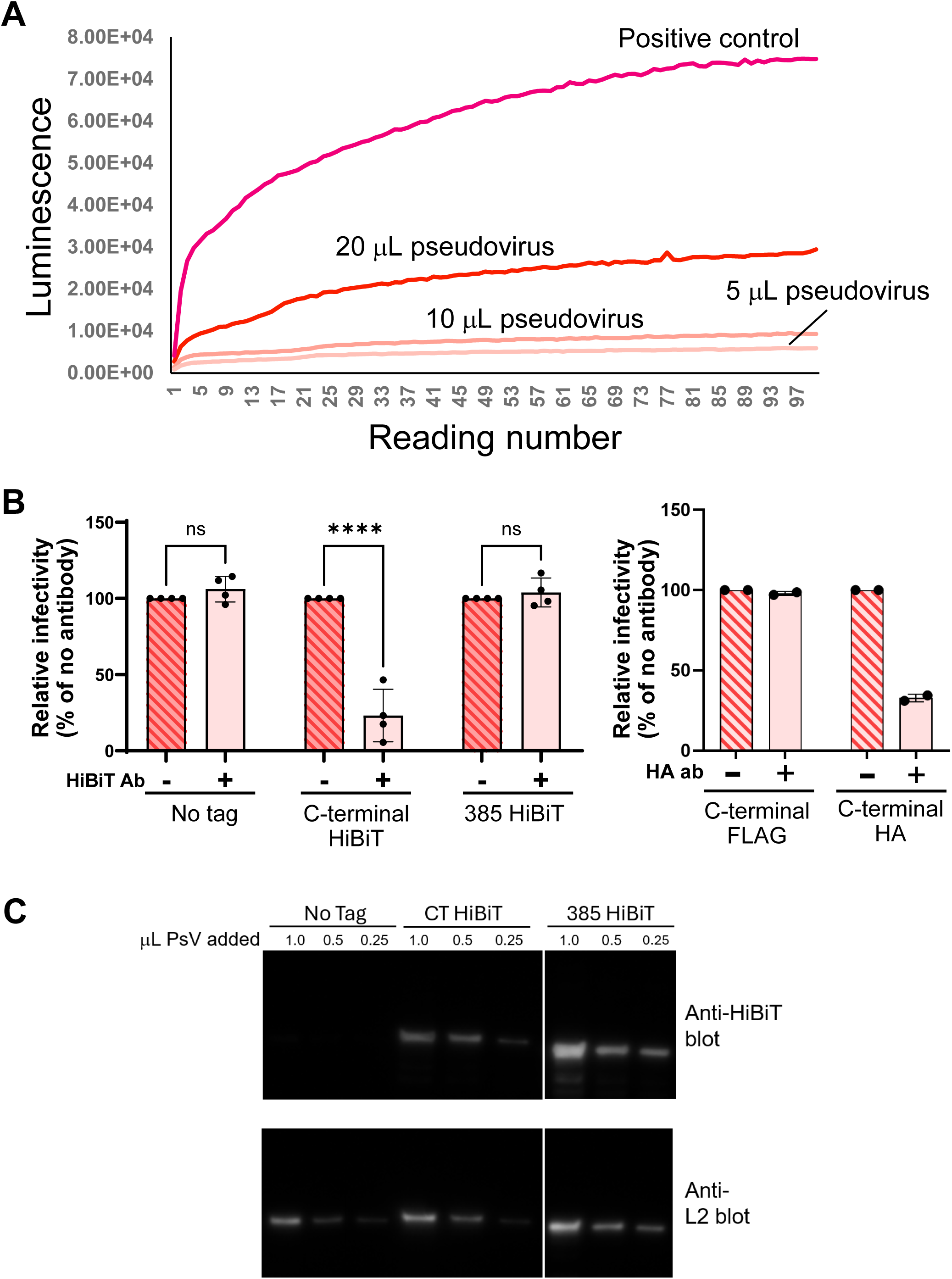
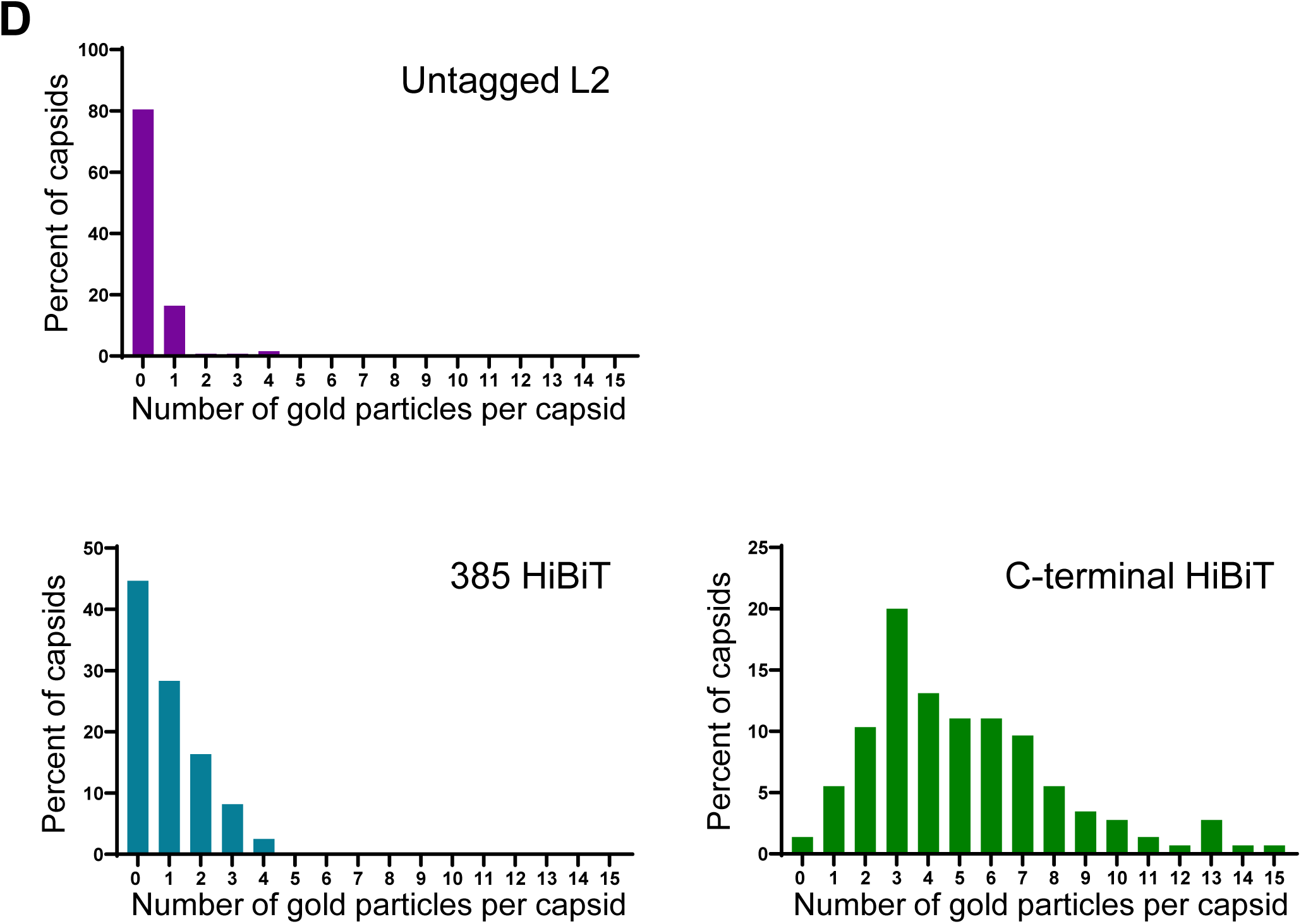
C-terminus of L2 is exposed on the surface of capsids. A. Dose dependence of luciferase reconstitution. Different amounts of a HPV16 PsV containing L2 with a C-terminal HiBiT tag were added to extracts containing LgBiT, and luciferase activity was measured in a luminometer at one-minute intervals. A L2-HiBiT fusion protein purified from bacteria was used as a positive control. B. Neutralization of HPV infection. Left, PsV containing untagged L2 or L2 with a HiBiT tag at position 385 or the C-terminus were incubated in the absence of antibody or in the presence of anti-HiBiT antibody. HeLa S3 cells were then infected and HcRed mean fluorescence intensity (MFI) was measured after 48 h. The MFI of each antibody-treated sample was normalized relative to its mock antibody-neutralized assay, with four biological replicates of the infection. Statistical significance was assessed as in Fig. 5C. ns, not significant; ****, p < 0.0001. Right, similar to left panel, but with PsV containing a FLAG tag or an HA tag at the C-terminus of L2, in the presence or absence of anti-HA antibody. C. Western blot of L2 containing a HiBiT tag at two different positions. HPV16 PsV were prepared containing no tag or a HiBiT tag at L2 position 385 or the C-terminus. The indicated amounts of each PsV preparation were subjected to SDS-polyacrylamide electrophoresis followed by western blotting with an antibody that recognizes HiBiT (top panel) or L2 (bottom panel). Irrelevant lanes were removed from the image of the gel. D. Distribution of gold tags on HPV PsV capsids. Graphs show histograms of the number of gold tags per capsid in PsV containing untagged L2, L2 containing a HiBiT tag at position 385, or L2 containing a C-terminal HiBiT tag. Samples were stained as in Figure 6C. Results shown are for biological replicate #2 in Figure 6B.

